# Substitutions in RNA-binding protein Hrp1 map a potential interaction surface with the yeast RNA polymerase II elongation complex

**DOI:** 10.1101/2025.05.07.652672

**Authors:** Moyao Wang, Payal Arora, Craig D. Kaplan, David A. Brow

**Affiliations:** Department of Biomolecular Chemistry, University of Wisconsin School of Medicine and Public Health, Madison, WI; Department of Biological Sciences, University of Pittsburgh, Pittsburgh, PA

## Abstract

Anti-termination factors for eukaryotic RNA polymerase II (RNAP II) that are released upon binding sequences in the terminator of nascent transcripts were proposed almost 40 years ago but few candidates have been found. Here we report genetic evidence that the yeast nuclear RNA-binding protein Hrp1, also known as Nab4 and CF1B, acts as an RNAP II anti-termination factor. A Lys to Glu substitution at residue 9 (K9E) of the Rpb3 subunit of RNAP II causes readthrough of Nrd1-Nab3-Sen1-dependent (NNS) terminators and cold-sensitive growth, as does Asp but not Ala, Met, Arg, or Gln substitution. These allele-specific phenotypes and the location of Rpb3-K9 suggests substitution with Glu or Asp stabilizes binding of an anti-termination factor via a salt bridge. A genome-wide selection for suppressors of the cold-sensitivity of Rpb3-K9E yielded an Arg to Gly substitution at residue 317 of Hrp1 in RNA recognition motif 2 (RRM2), consistent with the hypothesis. A targeted selection for suppressors of Rpb3-K9E in *HRP1* yielded substitutions in RRMs 1 and 2, an essential Met- and Gln-rich region C-terminal to the RRMs, and other mutations. We propose that Hrp1 binds to the RNAP II elongation complex via these regions to promote elongation and, in the presence of Rpb3-K9E, is less rapidly released upon binding terminator sequences in the nascent transcript, resulting in terminator readthrough. The Rpb3-K9E-suppressor substitutions in Hrp1 likely weaken binding to the RNAP II elongation complex, compensating for Rpb3-K9E.

## INTRODUCTION

All pathways of regulation of RNA synthesis by eukaryotic RNA polymerase II (RNAP II) must ultimately impact the enzyme directly, whether recruiting, blocking, or removing it from the DNA template, or exerting more subtle allosteric control of its activity. Given that RNAP II must transcribe long genes with a high degree of processivity to make functional mRNAs, mechanisms for terminating transcription must be tightly controlled. Yet, they must also be robust to prevent disruption of expression of a gene by extended transcripts coming from neighboring genes, especially in gene-dense genomes such as that of the yeast *Saccharomyces cerevisiae*. These properties imply that transcription termination involves multiple, cooperative molecular mechanisms. While much is known about the cis-acting termination signals and trans-acting termination factors for RNAP II, less is known about the molecular events that occur at RNAP II to stop its processive polymerase activity.

Termination of transcription requires slowing or pausing RNA polymerase followed by its removal from the DNA template by disrupting the elongation complex. In *S. cerevisiae* there are at least two pathways by which this occurs for RNAP II: the cleavage and polyadenylation factor-dependent (CPA) pathway and the Nrd1, Nab3, and Sen1-dependent (NNS) pathway. The CPA pathway is used primarily to terminate pre-mRNAs while the NNS pathway is used primarily for short non-coding RNAs, although there is some functional overlap of the two pathways (Kuehner et al. 2011; Arndt and Reines 2015; Rodríguez-Molina et al. 2023). CPA and NNS termination differ mechanistically. The CPA pathway involves endonuclease cleavage of the transcript, creating an uncapped 5’ end that is a substrate for a 5’ exonuclease “torpedo”. In contrast, the NNS pathway does not involve endonuclease cleavage and may instead use the 5’ to 3’ translocase Sen1 as a “helicase torpedo” (Steinmetz and Brow 1996; Porrua and Libri 2013; Rengachari et al. 2025). Both pathways may use, in addition, allosteric mechanisms to slow down or pause RNAP II. One proposed mechanism of allosteric termination in mammalian cells is recruitment of an anti-termination factor to RNAP II near the transcription start site that is released on binding to specific sequences in the nascent transcript at the terminator (Logan et al. 1987). Recently, SCAF4 and SCAF8, homologs of yeast Nrd1, were identified as potential RNAP II anti-terminator proteins in humans (Gregesen et al. 2019).

The main substrates of the NNS termination pathway are short, stable RNAP II transcripts, such as small nuclear (sn) and small nucleolar (sno) RNAs, and cryptic unstable RNAP II transcripts (CUTs), which include attenuated pre-mRNAs. NNS-terminated transcripts are targeted by the TRAMP complex for nuclear exosome 3’-processing (snRNAs and snoRNAs) or degradation (CUTs) (LaCava et al. 2005; Vasiljeva and Buratowski 2006; Tudek et al. 2014; Kim et al. 2016). Genome-wide selections and screens for trans-acting mutations that cause readthrough of snoRNA and CUT terminators identified genes encoding two RNA-binding proteins (Nrd1 and Nab3), an RNA/DNA helicase (Sen1), a protein phosphatase that acts on the Rpb1 CTD (Ssu72), the Rpb11 subunit of RNAP II, the polyA polymerase (Pap2/Trf4) from the TRAMP complex, and the Pcf11 subunit of pre-mRNA cleavage factor 1A (Steinmetz and Brow 1996; Steinmetz and Brow 2003; Steinmetz et al. 2006a; Loya et al. 2012). Substitutions in the cleavage and polyadenylation factor Hrp1 (Kessler et al. 1997; Minvielle-Sebastia et al. 1998), known also as Nab4 and CF1B, were also shown to cause readthrough of a subset of NNS terminators, including the attenuator of its own mRNA and the terminator of snoRNA snR82 (Steinmetz et al. 2006b; Kuehner and Brow 2008; Chen et al. 2017; Goguen and Brow 2023).

The substitution in Rpb11 that elicits readthrough of NNS terminators, Glu 108 to Gly (E108G), lies opposite the DNA entry site in RNAP II and about 4 nm from the binding site of the Sen1 helicase domain to a minimal RNAP II elongation complex in vitro (Rengachari et al. 2025; Figure 1). It may define a site of interaction with factors that influence termination. Rpb11 forms a heterodimer with the Rpb3 subunit of RNAP II and a targeted selection for dominant NNS terminator readthrough mutations in *RPB3* yielded a Lys to Glu substitution at residue 9 (K9E, Steinmetz et al. 2006a), which is immediately adjacent to Rpb11-E108 on the surface of the RNAP II elongation complex (Figure 1). The Rpb11-E108G substitution results in cold-sensitive growth, suggesting it stabilizes a transient interaction. Since the substitution elicits terminator readthrough, it could possibly stabilize binding of an anti-termination factor to the RNAP II elongation complex.

**Figure 1.**
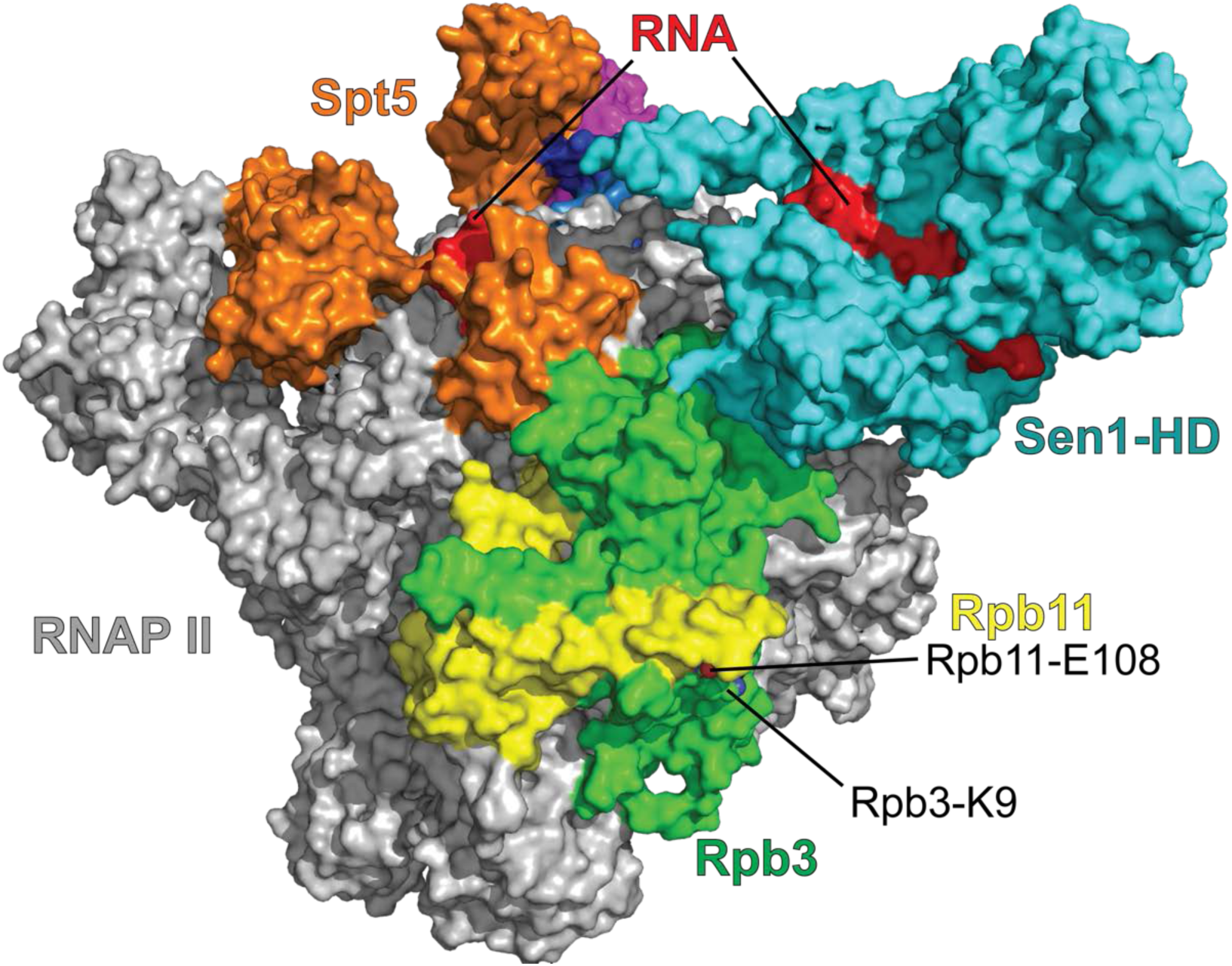
Sites of NNS readthrough substitutions in RNAP II subunits Rpb11 and Rpb3 are near a binding site of the Sen1 helicase domain. A cryo-EM structure of a minimal yeast RNA polymerase II elongation complex bound to purified Sen1 is from Rengachari et al. 2025 (PDB: 8rap). Residues in Rpb11 (yellow) and Rpb3 (green) that harbor substitutions causing readthrough of NNS terminators (Steinmetz *et al*. 2006a) are indicated. The nascent RNA is not resolved between the RNAP II exit site and the active site of the Sen1 helicase domain (Sen1-HD).

Here we show that the Rpb3-K9E substitution is also cold-sensitive. Of a range of substitutions tested in Rpb3-K9, only charge reversal to Asp or Glu result in cold-sensitivity and NNS terminator readthrough. A genome-wide selection for suppressors of the cold-sensitivity of *rpb3-K9E* yielded a substitution on the surface of RNA recognition motif 2 (RRM2) of Hrp1, Arg 317 to Gly (R317G), that also suppresses readthrough of two NNS terminators by the *rpb3-K9E* substitution. Thus, Hrp1-R317 is a candidate for forming a salt bridge with Rpb3-E9. A targeted selection for substitutions in *HRP1* that suppress the cold-sensitivity of *rpb3-K9E* yielded substitutions in the two RRMs, as well as a more C-terminal region rich in methionine and glutamine residues and next to a PY-NLS sequence at the C-terminus. Nonsense mutations were also obtained in the N-terminal low complexity domain and overexpression of the corresponding truncated proteins resulted in dominant suppression of *rpb3-K9E* cold-sensitive growth. Our results provide genetic evidence that Hrp1 is a general anti-termination factor for yeast RNAP II and suggest that it’s release from the elongation complex upon specific binding to the nascent transcript enhances termination.

## RESULTS

### Substitutions in Rpb3-K9 have allele-specific effects on growth and NNS termination consistent with a gain-of-interaction mechanism

To explore the mechanism of NNS terminator readthrough caused by *rpb3-K9E* we constructed an *RPB3* disruption strain (EJS300, Supplementary Table S1) and tested the recessive phenotypes of *RPB3-K9* alleles by plasmid shuffle. Like the *rpb11-E108G* substitution (Steinmetz et al. 2006a), the *rpb3-K9E* substitution is strongly cold-sensitive (Figure 2a). To test if the cold-sensitivity may be due to stabilizing a dynamic interaction, we made additional substitutions in Rpb3-K9, including alanine (A), aspartate (D), methionine (M), arginine (R), and glutamine (Q). Only the K9D substitution resulted in noticeable cold-sensitivity, which was weaker than that of K9E (Figure 2a). This result is consistent with the cold-sensitivity being due to stabilization of the interaction of residue 9 of Rpb3 with a positively charged amino acid. The weaker effect of the K9D substitution may be due to its shorter side chain, constraining interaction with another residue.

**Figure 2.**
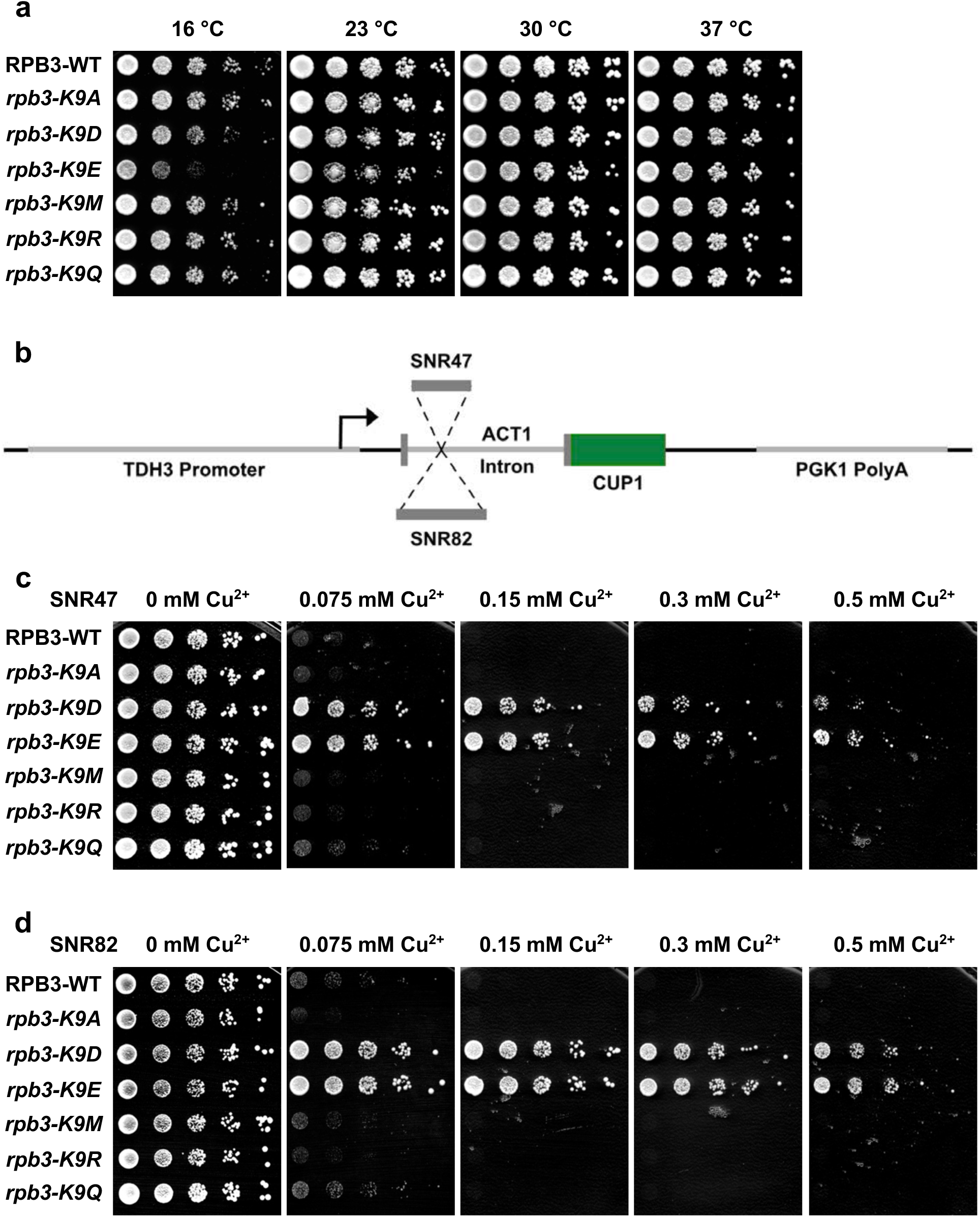
*rpb3-K9D* and *-K9E* substitutions confer cold-sensitivity and strong readthrough of the *SNR47* and *SNR82* terminators. **a**) Serial 8-fold dilutions of a haploid *11rpb3* strain containing the indicated alleles of *RPB3* on a centromere plasmid were spotted on YEPD plates and incubated at the indicated temperatures for two (30, 37 °C), three (23 °C), or nine (16 °C) days. **b**) Schematic of the ACT1-CUP1 fusion reporter (Lesser & Guthrie 1993) with either the *SNR47* or *SNR82* terminator region inserted in the *ACT1* intron. Serial 8-fold dilutions of haploid strains containing the indicated alleles of *RPB3* and the (**c**) *SNR47* or (**d**) *SNR82* ACT1-CUP1 reporter on separate centromere plasmids were spotted on synthetic complete medium containing the indicated concentration of copper sulfate and incubated at 30 °C for 4 days. Biological replicates are shown in Supplementary Figure S1.

To test for defects of the *RPB3* alleles in NNS termination, we assessed readthrough of the *SNR47* and *SNR82* snoRNA terminators when present in the intron of the ACT-CUP reporter gene (Figure 2b). With this reporter, strong readthrough confers strong copper resistance. Only the *rpb3-K9E* and *-K9D* alleles confer strong readthrough of these reporters (Figure 2c, d), with aspartate having a slightly weaker effect than glutamate. Thus, the cold-sensitivity correlates with weakening of NNS termination at 30 °C. We hypothesize that the *rpb3-K9D/E* substitutions stabilize an interaction between RNAP II and an anti-termination factor that normally is released from the elongation complex when a terminator sequence emerges from the polymerase.

### A selection for suppressors of *rpb3-K9E* identified a substitution in Hrp1

We reasoned that if the cold-sensitivity of the *rpb3-K9E* substitution is due to a stabilized interaction with a protein or nucleic acid, we should be able to isolate a destabilizing mutation in the interacting partner as a suppressor of the cold-sensitivity. We therefore isolated spontaneous suppressors of the 16 °C slow growth of this allele. Of the 20 colonies picked after 58-78 days of incubation at 16° C, all but two were white, one was pink, and one was red (Figure 3, Table 1). The red color comes from the accumulation of an oxidized precursor of adenine due to the *ade2-1* marker in the parent strain (Ugolini and Bruschi 1996 and references therein). The absence or decrease of red pigment could indicate phenotypic reversion of this marker.

**Figure 3.**
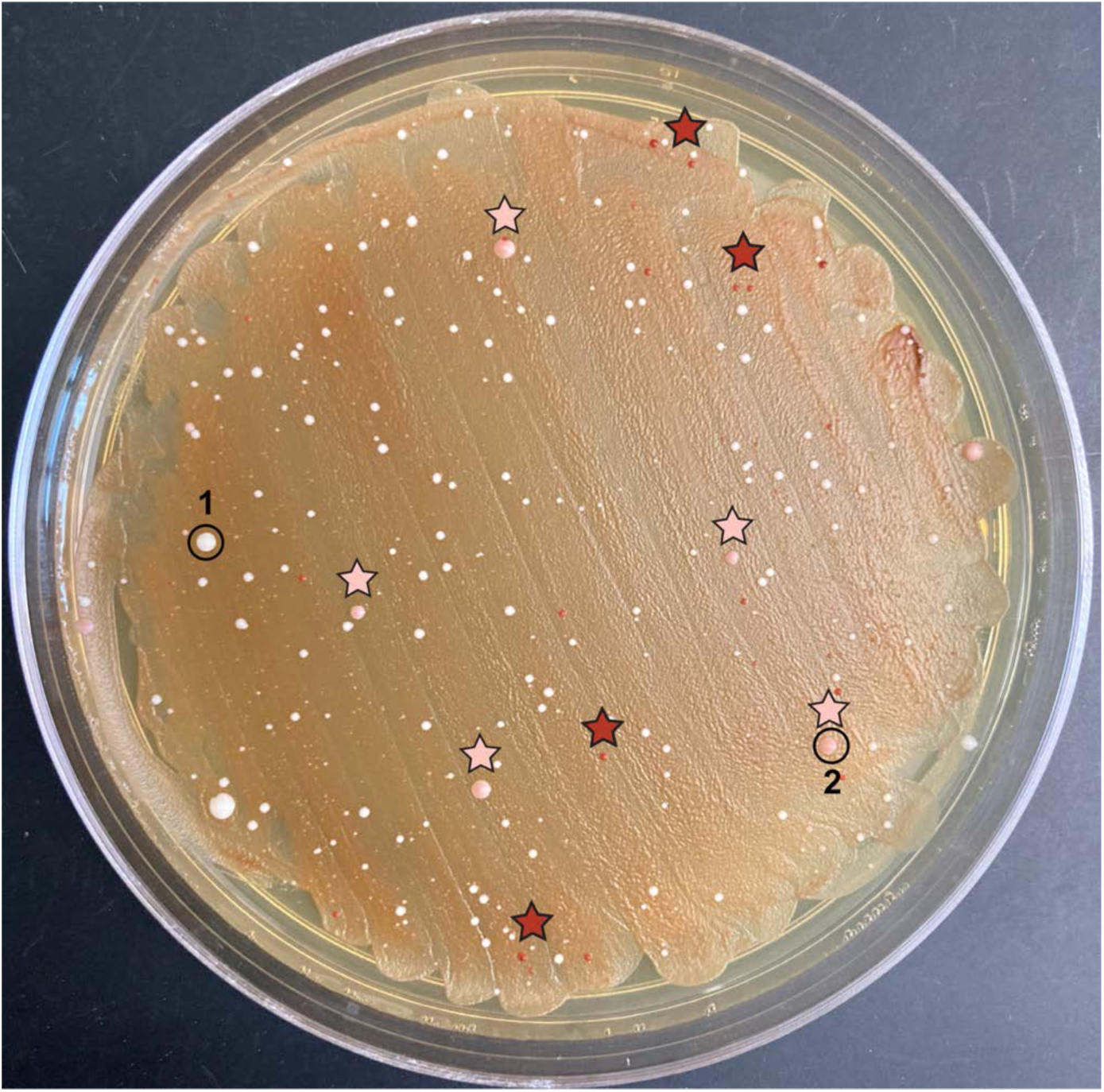
Plate B8 from the *rpb3-K9E*-suppressor selection after 11 weeks of growth at 16°. Apparent are white colonies of various sizes, medium-sized pink colonies (some of which are marked with pink stars), and tiny red colonies (some of which are marked with red stars). The colonies picked for genomic analysis, B8-1 and B8-2, are circled.

**Table 1.** Mutations identified in *rpb3-K9E*-suppressor strains.

| Clone | Days at 16° | Strain # | Colony color | Identified mutation(s) |
| --- | --- | --- | --- | --- |
| A10-1 | 58 | DAB177 | white | *SUP16/17/19* r.35G>T |
| A10-2 | 58 | DAB178 | white | *ade2-1* p.64X>L |
| B01-1 | 58 | DAB179 | white | *ade2-1* p.64X>Y |
| B01-2 | 58 | DAB180 | white | *ade2-1* p.64X>Y |
| B03-1 | 58 | DAB181 | white | *SUP16/17/19* r.35G>T |
| B06-1 | 58 | DAB182 | white | *ade2-1* p.64X>Y |
| B07-1 | 58 | DAB183 | white | *ade2-1* p.64X>Q |
| B10-1 | 58 | DAB184 | white | *ade2-1* p.64X>Q |
| A03-1 | 78 | DAB185 | red, clumpy |  |
| A04-1 | 78 | DAB186 | white | *SUP16/17/19* r.35G>T |
| A05-1 | 78 | DAB187 | white | *SUP16/17/19* r.35G>T |
| A06-1 | 78 | DAB188 | white | *ade2-1* p.64X>Y |
| A07-1 | 78 | DAB189 | white | *SUP16/17/19* r.35G>T |
| A09-1 | 78 | DAB190 | white | *SUP16/17/19* r.35G>T |
| B02-1 | 78 | DAB191 | white |  |
| B04-1 | 78 | DAB192 | white | *ade2-1* p.64X>Y |
| B05-1 | 78 | DAB193 | white | *ade2-1* p.64X>E |
| B06-2 | 78 | DAB194 | white | *SUP16/17/19* r.35G>T |
| B08-1 | 78 | DAB195 | white | *SUP16/17/19* r.35G>T |
| B08-2 | 78 | DAB196 | pink | *ADE4* p.90Q>E, *HRP1* p.317R>G |

To identify mutations responsible for suppressing *rpb3-K9E*, we screened the 20 strains using a custom targeted amplicon sequencing panel that probes 310 yeast genes involved in transcription (Supplementary Table S2). Only one suppressor strain has a mutation in the 310 genes: B8-2 has an Hrp1-R317G substitution. R317 is a surface exposed residue in RRM2 of Hrp1 (Figure 5c) and thus could mediate a direct contact with another factor that is modulated by RNA-binding. Hrp1-R317G was previously identified as causing weak readthrough of the Hrp1-dependent *SNR82* snoRNA terminator in the presence of wild-type RNAP II (Goguen and Brow 2003), which can be rationalized by relieving charge repulsion between Hrp1-R317 and Rpb3-K9, favoring extended occupancy of Hrp1 on RNAP II. Conversely, in the presence of Rpb3-K9E, Hrp1-R317G could break a Glu-Arg salt bridge, weakening an interaction between Hrp1 and Rpb3 and decreasing terminator readthrough.

To look for mutations outside of the 310 genes in the custom panel, we sequenced the whole genomes of all 20 cold-resistant strains. Surprisingly, nine of the strains have substitutions in the ochre (UAA) nonsense codon primarily responsible for loss of function of the *ade2-1* allele, changing it to a sense codon. An additional eight strains contain a substitution in the middle position of the anticodon of the *SUP16*, *SUP17*, or *SUP19* ochre-suppressor tRNA^Ser^ (Ono et al. 1981; Waldron et al. 1981) that should increase pairing with the UAA stop codon (U**G**A>U**U**A; Table 1). Which of these three paralogous *SUP* loci the mutant alleles came from could not be determined from the short reads, roughly a third of which were mutant. From these results we infer that the cold-sensitive growth phenotype of the *rpb3-K9E* substitution is a synthetic phenotype with the *ade2-1* mutation. This result also can explain why these suppressor strains are white, since they presumably have sufficient Ade2 activity to prevent accumulation of the red intermediate. In two strains, A3-1 and B2-1, a likely causative mutation could not be identified. A3-1 is red and presumably lacks an ochre suppressor, while B2-1 is white.

B8-2, which has the Hrp1-R317G substitution and is pink in color, also has a missense substitution in the *ADE4* gene (Table 1). The Ade4 enzyme, PRPP amidotransferase, catalyzes the first step of purine biosynthesis, five steps prior to the function of Ade2. It is not known how the *ade4-Q90E* substitution in this strain alters Ade4 function, although the light pink color of the colony suggests decreased flux through the adenine biosynthesis pathway, given the unaltered *ade2-1* allele and absence of a suppressor tRNA in this strain. It is known that mutation of the *ADE8* gene, which also encodes an enzyme that functions upstream of Ade2, reduces red pigment and imparts a strong growth advantage in the presence of *ade2-1* (Ugolini and Bruschi 1996). This result and the larger average colony size of the pink vs. red strains (Figure 3) suggest that the *ade4-Q90E* substitution enhanced the growth rate of the *hrp1-R317G* mutant.

### Different substitutions of Hrp1-R317 suppress Rpb3-K9E cold-sensitive growth and readthrough of NNS terminators, consistent with a loss-of-interaction mechanism

To test if the *hrp1-R317G* substitution is sufficient for suppression of the cold-sensitivity of *rpb3-K9E*, we introduced the *hrp1-R317G* allele into an *RPB3/HRP1* double disruption strain that contains the *rpb3-K9E* substitution on a centromere plasmid. Since we hypothesized that *hrp1-R317G* suppresses the cold-sensitive growth of *rpb3-K9E* by disrupting a salt bridge between *hrp1-R317* and *rpb3-K9E*, we also tested the *hrp1-R317E* substitution. This substitution would reinstate the repulsive interaction predicted in the wild-type strain between *hrp1-R317* and *rpb3-K9* and therefore may suppress the cold-sensitive phenotype as well as, or better than, *hrp1-R317G*. In addition, we introduced *hrp1-R317Q* substitution to test whether a polar but noncharged amino acid (glutamine) could suppress the cold-sensitive phenotype.

We found that the *rpb3-K9E* substitution alone confers slower growth at all temperatures tested in the double disruption strain (Figure 4a, row 5). The reason for the temperature-independent slow growth in this strain background is unknown. Nevertheless, the strain still exhibits strong cold-sensitivity at 16 °C and moderate cold-sensitivity at 23 °C. Importantly, the *hrp1-R317G* substitution strongly rescues both the cold-sensitive and slow-growth phenotypes of *rpb3-K9E* (Figure 4a, row 6), indicating that the suppression observed in the whole-genome selection strain B8-2 is largely or completely due to the *hrp1-R317G* substitution. The *hrp1-R317Q* substitution exhibits weaker suppression at all temperatures, perhaps because the Gln side chain can form a hydrogen bond with Rpb3-E9. The *hrp1-R317E* substitution exhibits slightly weaker suppression of *rpb3-K9E* cold-sensitivity than R317G, possibly due to other effects of the substitution. Interestingly, a potential reversed salt bridge from the Rpb3-K9/Hrp1-R317E combination (Figure 4a, row 4) does not result in cold-sensitivity. Perhaps the adjacent Rpb11-E108 residue prevents the salt bridge from being stable in this orientation.

**Figure 4.**
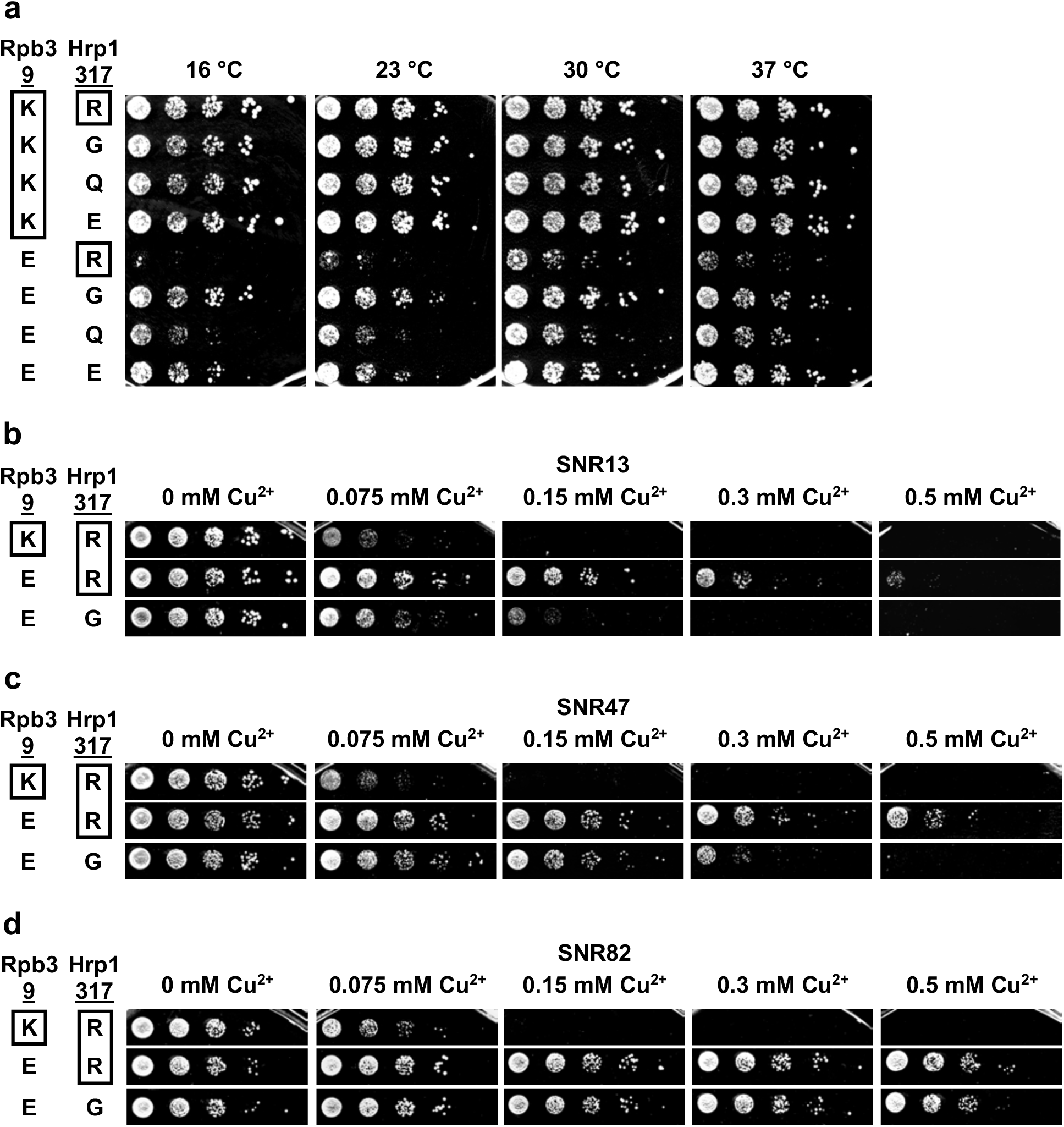
*hrp1-R317* substitutions exhibit allele-specificity for suppression of *rpb3-K9E* cold-sensitivity and terminator-specific readthrough suppression. **a)** Serial dilutions of haploid strains containing the indicated alleles of *RPB3* and *HRP1* (wildtype residues are boxed) on separate centromere plasmids were spotted onto YEPD plates and incubated at the indicated temperatures for 2 (30, 37 °C), 3 (23 °C), and 7 (16 °C) days. **b-d**) Serial dilutions of haploid strains containing the indicated alleles of *RPB3* on a centromere plasmid with indicated alleles of *HRP1* in the genome on CSM -Leu medium containing the indicated concentrations of copper sulfate were incubated at 30 °C for 4 days. All rows for each panel came from the same plate. Biological replicates are shown in Supplementary Figure S2.

We next tested the ability of the *hrp1-R317G* substitution to suppress the *rpb3-K9E* terminator readthrough phenotype. We first introduced the R317G substitution into the *HRP1* genomic locus of an *RPB3* disruption strain containing the rpb3-K9E substitution on a centromere plasmid. We then assessed the readthrough of several NNS terminators using the ACT1-CUP1 reporter gene containing different snoRNA terminator sequences in its intron (Figure 2b). The *hrp1-R317G* substitution moderately suppressed *SNR13* readthrough, weakly suppressed *SNR47* readthrough, and did not suppress *SNR82* readthrough caused by Rpb3-K9E (Figure 4b-d). Hrp1-R317G causes weak readthrough of *SNR82* in a wild-type Rpb3 background (Goguen and Brow, 2023), which may be masking weak suppression of *SNR82* readthrough in the presence of Rpb3-K9E. Nevertheless, the partial suppression by Hrp1-R317G of *SNR13* and *SNR47* readthrough by Rpb3-K9E supports the hypothesis that the cold-sensitivity is due to NNS terminator readthrough.

### A selection for mutations in *HRP1* that suppress *rpb3-K9E* cold-sensitivity defines potential interaction surfaces

To identify other residues in *HRP1* that may interact with RNAP II or its elongation factors, we selected suppressors of the cold-sensitive growth of *rpb3-K9E* after mutagenizing the *HRP1* ORF by amplification with Taq DNA polymerase and co-transforming it with a gapped centromere plasmid. After selection for the recombined *HRP1* plasmid, the transformation plates were replica plated onto SC 5-FOA medium to select for the loss of the *URA3*-marked wildtype *HRP1* plasmid. Surviving colonies were replica plated to YEPD and incubated at 16 °C to select for the suppression of the cold-sensitive growth phenotype of *rpb3-K9E*. Eight colonies were picked after 5 days and twenty more colonies were picked after 6 days (Supplementary Table S2). The much shorter growth time required compared to the spontaneous suppressor selection is likely due to prior mutagenesis and the selection for growth of colonies rather than single cells at 16 °C.

To identify potential suppressor mutations, the *HRP1* plasmid was rescued from each of the 28 yeast strains and sequenced. Plasmids from 14 of the strains have nonsynonymous mutations in *HRP1*. Ten strains have no mutation in the *HRP1* ORF and two have only synonymous mutations. Some of these 12 strains may have a spontaneous suppressor elsewhere in the genome. Indeed, two of them form white colonies, suggesting they have mutated or suppressed the nonsense codon in the *ade2-1* allele, as occurred in the genome-wide selection. We were unable to rescue a plasmid from two of the strains.

Four of the 14 strains with nonsynonymous DNA substitutions in *HRP1* have early nonsense mutations: Q29X, Q41X (obtained twice), and Q130X (Figure 5a, Supplementary Figure S3). As these nonsense codons are in the sole copy of *HRP1* and are well upstream of essential domains of Hrp1 (Goguen and Brow 2023), they would normally be expected to be lethal. Shuffling these plasmids into a naïve *rpb3-K9E* strain confirmed their viability and suppressor activity, comparable to the R317G substitution (Figure 5d, rows 6, 12, and 13). We previously selected viable nonsense mutations in *NRD1* (Steinmetz and Brow 1996) and a frameshift mutation in *HRP1* (Goguen and Brow 2023). Because these genes are negatively autoregulated by NNS attenuators (Steinmetz et al. 2001, Steinmetz et al. 2006b), their mRNA levels can increase more than 10-fold to allow sufficient stop codon readthrough or frameshifting to produce nearly normal levels of full-length protein. To confirm that is the case for these *HRP1* mutations, we inserted N-terminal GFP tags into wild-type and nonsense containing *HRP1* alleles and compared their protein production by Western blot (Figure 5b). As predicted, the nonsense alleles of *HRP1* produce at least half the normal amount of full-length protein in addition to 2- to 3-fold greater amounts of truncated proteins of the expected mobility given the location of the nonsense codons. Thus, either the truncated proteins or full-length protein with a substitution at the nonsense codon (or both) are responsible for suppression of the *rpb3-K9E* substitution (see next section). Since the stop codons created by mutation of these glutamine codons are UAA (Q29X, Q41X) and UAG (Q130X), readthrough is expected to insert tyrosine or glutamine at these positions (Blanchet et al. 2014).

**Figure 5.**
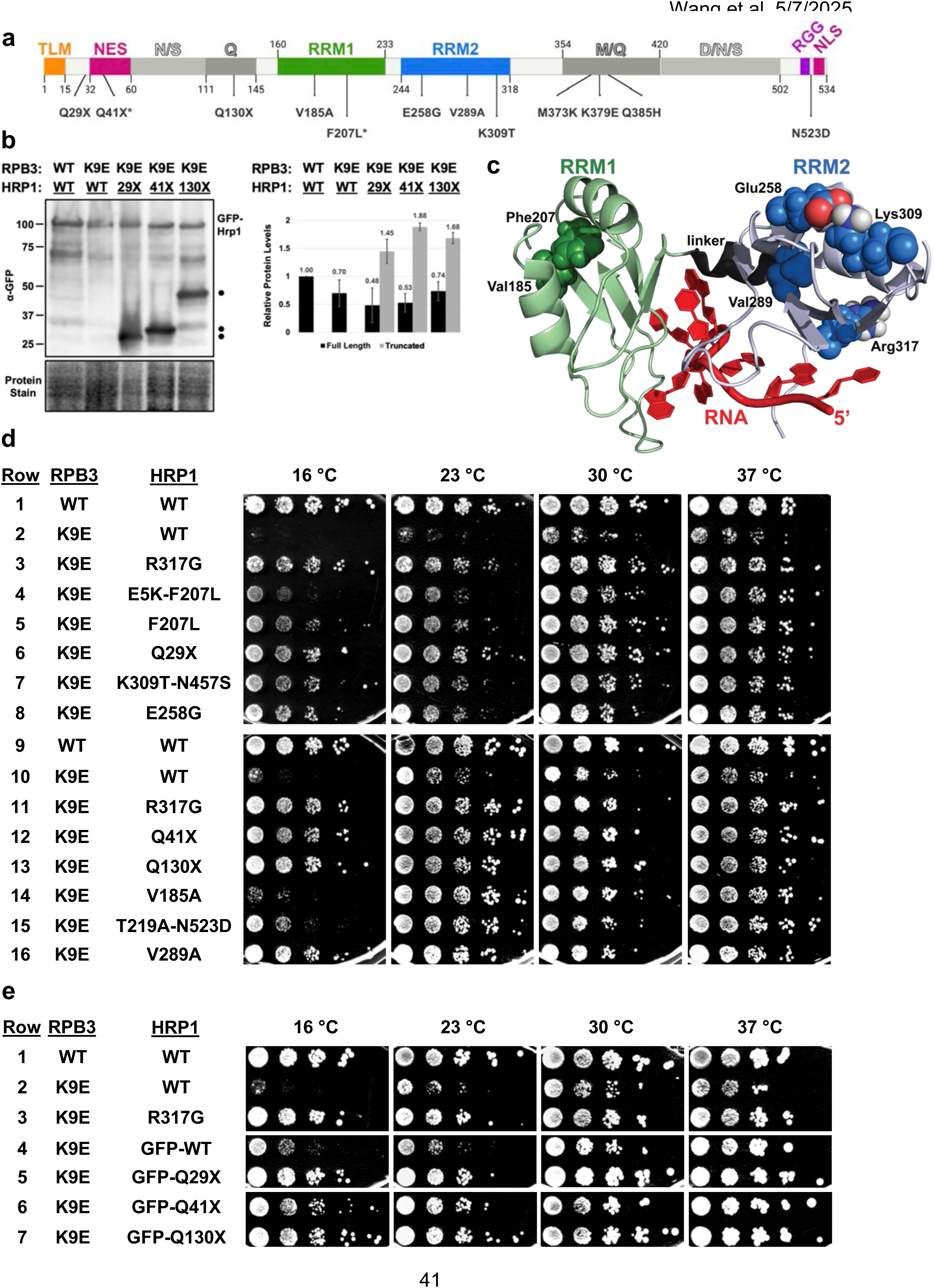
Mutations identified in the *HRP*1-targeted selection for suppressors of *rpb3-K9E* cold-sensitive growth. **a**) Mutations that confer cold-resistance to an *rpb3-K9E* strain mapped on the primary structure of Hrp1. Amino acid residue numbers are shown. TLM, TIM-like motif; NES, nuclear export sequence; N/S, Asn/Ser-rich low complexity domain (LCD); Q, Gln-rich LCD; RRM, RNA recognition motif; M/Q, Met/Gln-rich LCD; D/N/S, Asp/Asn/Ser-rich LCD; RGG, two Arg-Gly-Gly repeats; PY-NLS, proline-tyrosine nuclear localization signal. Asterisks indicate mutations obtained twice. **b**) Anti-GFP Western blot of cell extracts from strains with N-terminal GFP-tagged wildtype and nonsense *HRP1* alleles. The position of full-length GFP-HRP1 is indicated and prematurely terminated proteins are marked with dots. The average and range of relative protein levels from biological replicate experiments is shown at right. **c**) Locations of *rpb3-K9E*-suppressor substitutions in an NMR structure of Hrp1 RRMs 1 and 2 bound to (AU)_4_ (PDB: 2CJK; Pérez-Cañadillas 2006). **d,e**) Serial dilutions (1:8) of haploid strains containing the indicated alleles of *RPB3* and *HRP1* on separate centromere plasmids were spotted onto YEPD plates and incubated at the indicated temperatures for 2 (30, 37 °C), 3 (23 °C), or 9 (16 °C) days. Biological replicates are shown in Supplementary Figure S4.

Of the remaining ten alleles, seven have single amino acid substitutions and three have double substitutions. The single substitutions are in RRM1, RRM2, and an essential methionine- and glutamine-rich (M/Q) low complexity domain C-terminal to RRM2 (Figure 5a, Supplementary Figure S3). When mapped to an NMR structure of the Hrp1 RRMs bound to (UA)_4_ RNA (Pérez-Cañadillas 2006), the two single substitutions in RRM1 (V185A and F207L) are in residues that form a van der Waals interaction pair that anchors the beta sheet to the two alpha helices, distal to the RNA-binding face (Figure 5c). Suppressor mutations identified in both residues substitute a large hydrophobic residue with a smaller counterpart that could disrupt the van der Waals interaction and alter the fold and/or dynamics of RRM1. Both substitutions are weaker suppressors, especially V185A, but their suppressor activity is clearly seen at 23 °C (Figure 5d, rows 5 and 14).

The F207L substitution was also obtained in combination with E5K (Supplementary Table S2), but this double substitution is a weaker suppressor of Rpb3-K9E cold-sensitivity than F207L alone (Figure 5d, rows 4 and 5), suggesting that E5K counteracts suppression by F207L or is cold-sensitive itself. Indeed, combining Hrp1-E5K mutation with Rpb3-K9E leads to an even stronger cold sensitive phenotype compared to Hrp1-WT and Rpb3-K9E combination (Figure 6c, rows 4-6). Hrp1-E5 is in the highly acidic, conserved TIM-like motif (TLM), deletion of which induces NNS terminator readthrough (Goguen and Brow 2023). The charge switch substitution may disrupt an interaction with an unidentified protein.

**Figure 6.**
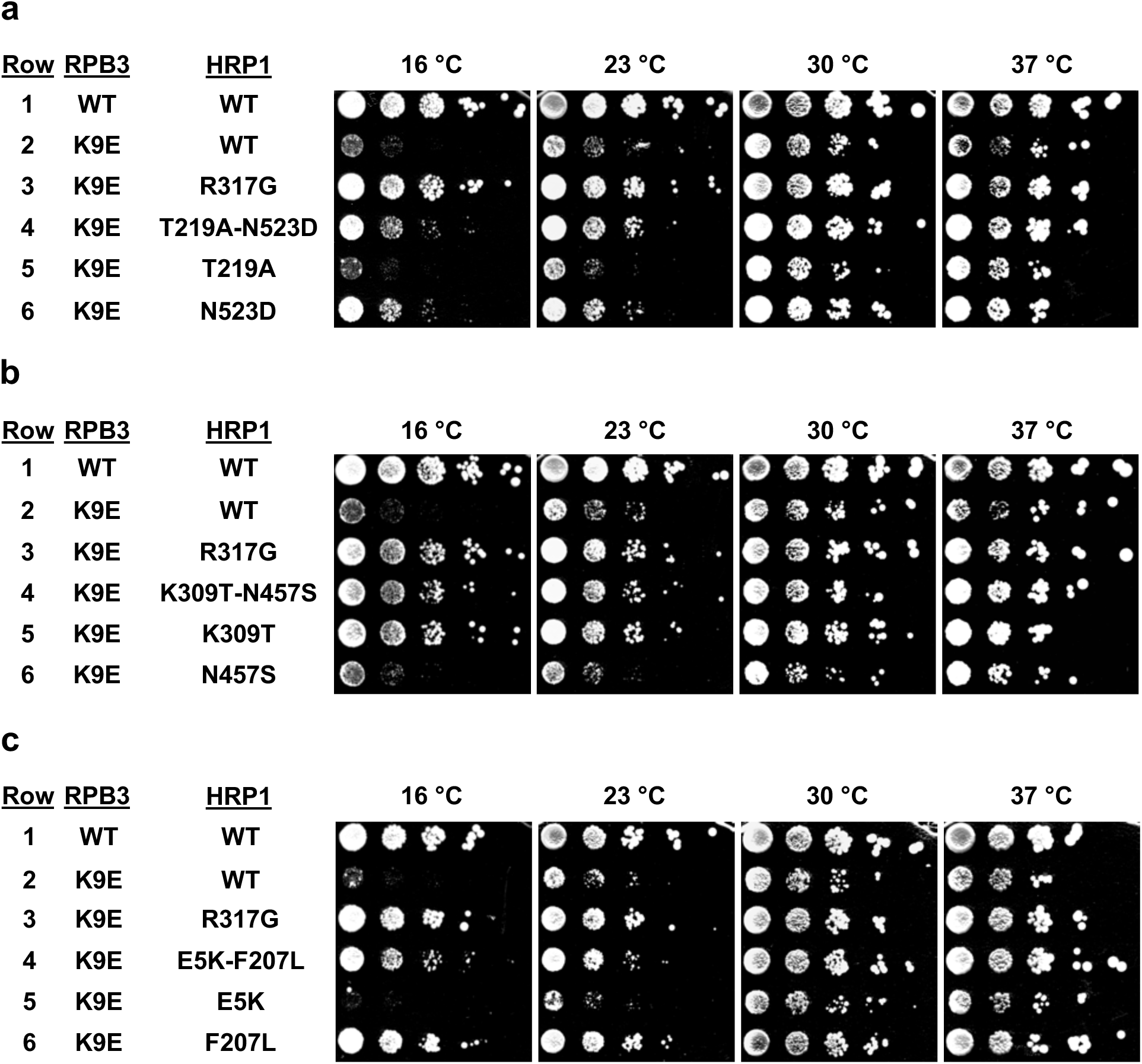
Each of the double substitutions selected in *HRP1* comprise one suppressor and one passenger mutation. Serial dilutions (1:8) of haploid strains containing the indicated alleles of *RPB3* and *HRP1* on separate centromere plasmids were spotted onto YEPD plates and incubated at the indicated temperatures for 2 (30, 37 °C), 3 (23 °C), and 9 (16 °C) days. The starting double mutant is in row 4 of each panel and each single mutant is in rows 5 and 6.

The other mutation identified in RRM1, T219A, is in combination with N523D, which lies between two RGG tripeptides and the C-terminal proline-tyrosine nuclear localization sequence (PY-NLS, Supplementary Figure S3) (Lange et al. 2008). This double mutation weakly suppresses the cold-sensitivity of *rpb3-K9E*, similar to the level of suppression seen with N523D alone (Figure 6a, rows 4-6). T219A alone exhibits no suppression of *rpb3-K9E*. Thus, N523D is responsible for suppression in the double mutant.

The two single substitutions in RRM2 are E258G and V289A. The only other substitution in RRM2 is K309T, which is combination with N457S in the D/N/S low complexity domain. However, N457S exhibits no suppression, while K309T alone is a strong suppressor, similar to R317G (Figure 6b, rows 3-6). Notably, E258 and K309 form a salt bridge on the surface of RRM2 opposite the RNA-binding face (Figure 5c). V289A, is in the beta sheet but faces away from the RNA, buried in the core fold of the RRM. None of the substitutions are adjacent to R317, the site of the founding suppressor substitution. Thus, like the substitutions in RRM1, the suppressor substitutions in RRM2 do not appear to directly influence RNA-binding but instead are in residues that stabilize the RRM fold.

The three remaining single substitutions are in the M/Q domain: M373K, K379E, and Q385H (Supplementary Table S2). AlphaFold (Jumper et al. 2021; Varadi et al. 2023) predicts two alpha-helices in the M/Q domain and the three substitutions are all on the longer of the two (Supplementary Figure S3, Figure 7a). M373K, K379E, and Q385H each moderately suppress the cold-sensitivity and combining two of them does not noticeably increase suppression (Figure 7b, rows 4-7). The alpha-helix also contains one of the four substitutions in *hrp1-7*, Y383H (Goguen and Brow 2023). This substitution only weakly suppresses the cold-sensitivity (Figure 7b, row 7). The clustering of these substitutions and the essential nature of the M/Q domain suggests that this alpha helix makes important contacts.

**Figure 7.**
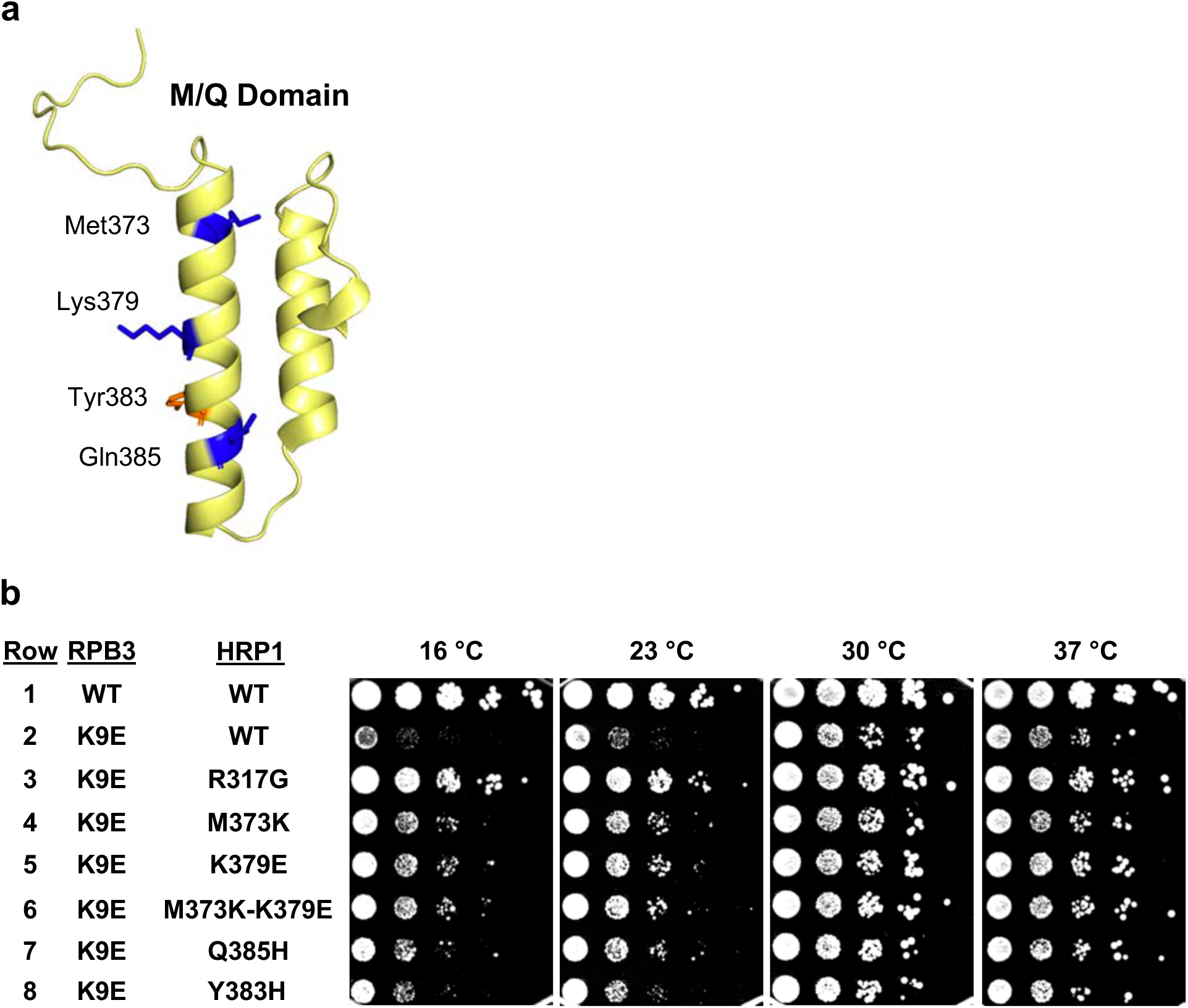
Hrp1 M/Q domain substitutions are weaker *rpb3-K9E*-suppressors than R317G, even in combination. **a**) AlphaFold-predicted structure of Hrp1 M/Q domain. **b**) Serial dilutions (1:8) of haploid strains containing the indicated alleles of *RPB3* and *HRP1* on separate centromere plasmids were spotted onto YEPD plates and incubated at the indicated temperatures for 2 (30, 37 °C), 3 (23 °C), and 9 (16 °C) days.

### Overexpression of residues 1-28 of Hrp1 suppresses rpb3-K9E cold-sensitive growth

To test if the truncated proteins created by the nonsense mutations in *HRP1* are responsible for suppression of *rpb3-K9E*, we expressed the corresponding truncated forms of Hrp1 from a high-copy plasmid in an *RPB3/HRP1* double disruption strain that contains both wildtype *HRP1* and *rpb3-K9E* alleles on a single low-copy plasmid. Transcription of the truncated *HRP1* alleles from the constitutive *TDH3* promoter and 5’-UTR on the high-copy pG1 plasmid prevented their repression by the *HRP1* attenuator. Overexpression of residues 1-28 of Hrp1 strongly suppresses the cold sensitivity of rpb3-K9E, while the overexpression of residues 1-40 of Hrp1 confers much weaker suppression (Figure 8a, Rows 4 and 5). We were unable to transform this strain with a high-copy plasmid expressing Hrp1(1-129), corresponding to the Q130X mutation, suggesting that high-level overexpression of the NES, N/S, and/or Q domains (Figure 5a) may be toxic. These results show that overexpression of the first 28 residues of Hrp1 is sufficient to suppress *rpb3-K9E* cold-sensitivity, presumably by competing for binding of this region to the RNAP II elongation complex.

**Figure 8.**
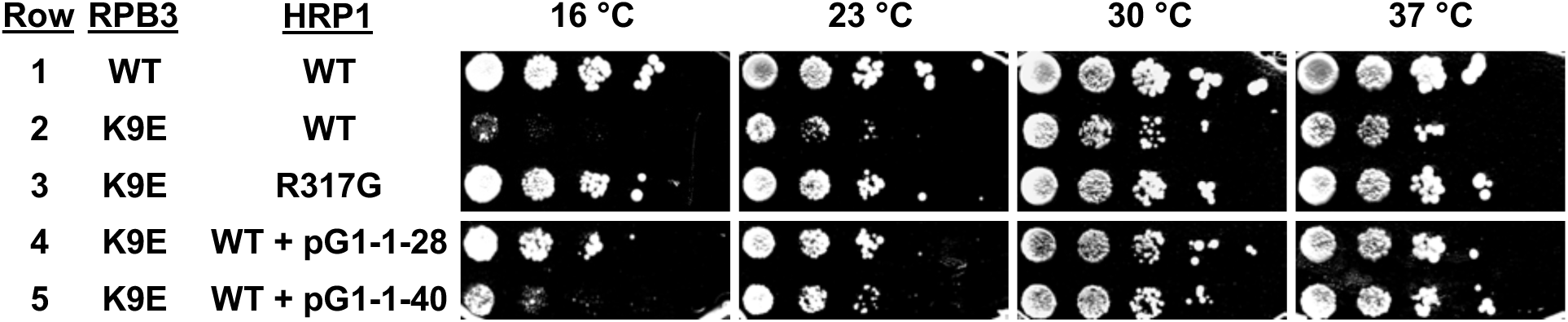
Overexpression of Hrp1-1-28 and 1-40 suppress Rpb3-K9E’s cold sensitivity. Serial dilutions (1:8) of haploid strains containing the indicated alleles of *RPB3* and *HRP1* on separate centromere plasmids (Row 1-3) or on the same centromere plasmid with Hrp1 overexpression construct (Row 4-5) were spotted onto YEPD plates and incubated at the indicated temperatures for 2 (30, 37 °C), 3 (23 °C), and 9 (16 °C) days.

### The Hrp1-N523D substitution does not disrupt nuclear localization of Hrp1

Since the Hrp1-N523D substitution is on the edge of the core PY-NLS sequences (Lange et al. 2008), we tested if it suppresses the cold sensitivity of Rpb3-K9E by altering the localization pattern of Hrp1. We transformed a N-terminal GFP-tagged HRP1 plasmid into a strain with integrated RFP-tagged nuclear marker NHP6A. We observed strong nuclear localization of both GFP-Hrp1 and GFP-Hrp1-N523D (Supplementary Figure S8), indicating that Hrp1-N523D substitution does not impact the nuclear localization of Hrp1.

## DISCUSSION

The fact that Hrp1 facilitates both the CPA and NNS RNAP II termination pathways suggests that it may be working by a more general mechanism than the pathway-specific factors. We propose that that Hrp1 acts as an anti-terminator protein that promotes RNAP II elongation but disengages from the elongation complex upon binding a high-affinity site in the terminator of the nascent transcript (Figure 9a). This model predicts that substitutions that weaken specific RNA binding promote terminator readthrough while substitutions that weaken binding to, or allosteric activation of, the elongation complex reduce terminator readthrough. Our prior selection for terminator readthrough mutations in *HRP1* (Goguen and Brow 2023) and the *rpb3-K9E*-suppressors substitutions reported here support this model, as the former localize primarily to the RNA-binding surfaces, while the latter are distal to the bound RNA on the periphery of the RRMs (Figure 9b).

**Figure 9.**
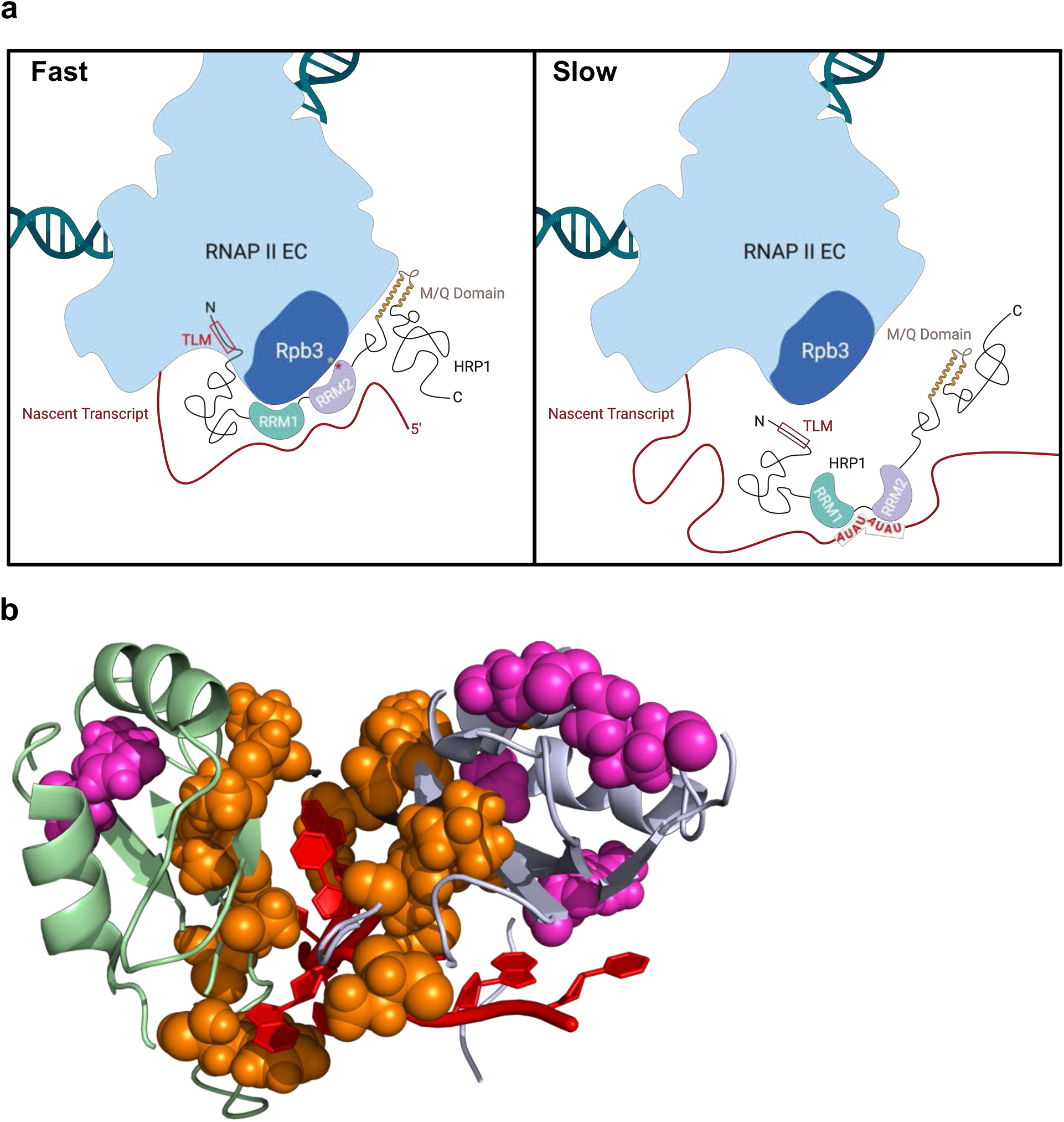
**a**) Model of Hrp1 interaction with the RNAP II elongation complex before (Fast) and after (Slow) binding to UA repeats in the nacent transcript. Gold and red stars represent the locations of Rpb3-K9 and Hrp1-R317, respectively. **b**) Locations of *SNR82* terminator readthrough substitutions (orange; Goguen and Brow 2023) and *rpb3-K9E*-suppressor substitutions (magenta; this study) in the RRMs of Hrp1. The structure (PDB: 2CJK; Pérez-Cañadillas 2006) is in the same orientation as in Figure 5c.

### *Rpb3-K9E*-suppressor mutations in the Hrp1 RRMs likely perturb their tertiary structures

The two *rpb3-K9E* suppressor mutations in RRM1 are each expected to perturb a van der Waals pair (V185-F207) that ends a beta strand 2/3 hairpin (Figure 5c). Both residues are perfectly conserved between five diverse fungal species (Supplementary Figure S9), suggesting this interaction is functionally important. The beta turn of this hairpin contains a proline-alanine dipeptide, P194/A195. The *hrp1-7* allele contains the substitution A195P, which would create a diproline in this turn. A195P alone exhibits no readthrough of the *HRP1* attenuator or *SNR82* terminator but instead appears to <u>decrease</u> expression of the HRP1-CUP1 reporter, suggesting it may decrease anti-terminator activity of Hrp1 (Goguen and Brow 2023). Thus, perturbation of the RRM1 beta strand 2/3 hairpin from either end may disrupt interaction of Hrp1 with the elongation complex.

We identified four *rpb3-K9E* suppressor mutations in RRM2: E258G, V289A, K309T, and R317G. E258 and K309 form a salt bridge connecting alpha helix 1 to the loop between alpha helix 2 and beta strand 4. R317 is at the end of beta strand 4, and V289 contacts F263 in alpha helix 1. All but V289 are surface exposed. Interestingly, V289A was also obtained as part of triple mutant with T186A (in RRM1) and Q282L (in RRM2) in a selection for *SNR82* terminator readthrough (allele 123 in Goguen and Brow 2023). T186 packs against V185 and F207, the *rpb3-K9E*-suppressors in RRM1, so both V289A and T186A are expected to diminish readthrough. In contrast, Q282 appears to hydrogen bond to the phosphoribose backbone of bound RNA and its substitution with leucine may be responsible for the terminator readthrough activity. As the triple mutant is viable, the V289A and T186A substitutions may offset an otherwise detrimental strong readthrough effect of Q282L, although this remains to be tested.

In summary, for the two RRMs of Hrp1, terminator readthrough substitutions are consistent with decreased recognition of the termination signal in the RNA, while *rpb3-K9E*-suppressor substitutions are consistent with weakening positive interactions with the RNAP II elongation complex. However, further experiments will be required to substantiate these conclusions.

### Genetic evidence for contacts between the Hrp1 N- and C-termini and the RNAP II elongation complex

We show that overexpression of the N-terminal 28 residues of Hrp1, either from a nonsense allele that produces both truncated and full-length protein or from a high-copy plasmid in the presence of wildtype Hrp1, suppresses the cold-sensitivity of *rpb3-K9E*. The first 15 residues of Hrp1 contains a sequence similar to the TFIIS interacting motif (TIM), which we call the TIM-like sequence or TLM (Goguen and Brow 2023). The TIM is common in proteins involved in RNAP II elongation (Cermakova et al. 2021). It is also similar to the Nrd1 interaction motif that binds the Nrd1 CTD-interacting domain (CID) and AlphaFold2 (Jumper et al. 2021) predicts the interaction of the N-terminus of Hrp1 with the Nrd1 CID. The TLM is conserved among fungi (Supplementary Figure S9) and the Hrp1-E5K substitution obtained in combination with the F207L *rpb3-K9E*-suppressor reverses the charge of this conserved acidic residue. It will be of interest to determine if this substitution reduces the ability of Hrp1(1-28) overexpression to suppress the cold-sensitivity of *rpb3-K9E*.

A curious property of the viable deletion of residues 2-15 of Hrp1, corresponding to the TLM, is that the level of Hrp1 decreases about 10-fold as measured by Western blot with Hrp1 IgG (Gougen and Brow, 2023). At the time we thought that the N-terminus of Hrp1 may be an important epitope for the polyclonal rabbit IgG, but in this study we found that the same anti-Hrp1 IgG is unable to detect the truncated Hrp1 produced from the nonsense alleles (Supplementary Figure S10), indicating that they don’t contain major epitopes. Thus, the decrease in Hrp1 levels appears to be real. Furthermore, an N-terminal GFP tag on Hrp1 decreases the protein level 3-fold (Goguen and Brow 2023). We also previously found that deletion of the Hrp1 D/N/S LCD, which is lethal, resulted in a 10-fold decrease in protein from both the deleted allele and the wildtype allele present to keep the cells alive (Goguen and Brow 2023). To explain this trans effect, we proposed that the D/N/S LCD is a negative autoregulatory domain that binds to the RRMs and competes for RNA-binding. Its deletion would therefore increase binding of Hrp1 to the *HRP1* attenuator, resulting in less mRNA. The same mechanism could explain the effect of deleting the TLM, if the TLM and D/N/S LCD are both required for negative regulation. The bacterial group 1 sigma factor N-terminal 1.1 domain provides a precedent for such negative regulation (Dombroski et al. 1993). This domain autoinhibits DNA binding by sigma factor in the absence of RNAP. When RNAP binds sigma factor the 1.1 domain moves from the sigma DNA-binding region to the RNAP DNA cleft, allowing binding of sigma to the promoter elements (Schwartz et al. 2008).

A similar activity of the Hrp1 TLM is consistent with its −7 net charge, or up to −10 if serines 2 and 3 and tyrosine 12 are phosphorylated, which they are found to be in several phosphoproteome studies (https://www.yeastgenome.org/locus/S000005483/protein). Binding of the TLM to the RRMs would presumably require the D/N/S LCD. If Hrp1 is regulated like group 1 sigmas, binding to RNAP II would release inhibition of the RRMs. Thus, deletion of the TLM would not increase attenuation of Hrp1 mRNA. Indeed, we found that Hrp1-ΔTLM mRNA increases 4-fold compared to wildtype Hrp1 mRNA (Goguen and Brow 2023), in contrast to the 10-fold reduction in protein, so the decrease in Hrp1 protein may be due to post-transcriptional steps of gene expression. Perhaps stronger binding of free Hrp1-ΔTLM to the 5’-UTR of its mRNA inhibits nuclear export and/or translation.

The M/Q-rich low complexity domain is known to be essential (Goguen and Brow 2023) but its function is unknown. The fact that a third of our missense suppressor substitutions map to a small region of this domain suggest the region is involved in termination. The same region of the M/Q LCD is the most highly conserved (Supplementary Figure S9). The methionine of the M373K substitution is perfectly conserved in five diverse fungi and the lysine of the K379E charge reversal substitution is a positively charged residue in four of the five fungal species. However, all three substitutions in the M/Q-rich LCD are fairly weak suppressors, so the interaction involving this region may not be as strong as that with the RRMs or TLM, or the substitutions may have other, negative consequences. The N523D substitution is also a weaker suppressor. It lies between the conserved, C-terminal PY nuclear localization signal and a conserved RGG sequence, in which the arginine is known to be methylated in *S. cerevisiae* (Xu and Henry 2004). Given that N523D does not noticeably affect the nuclear localization of Hrp1, it may alter an interaction with the RGG repeat.

### How does Rpb11-E108 interact with Rpb3-K9?

Both the Rpb11-E108G and Rpb3-K9E substitutions confer cold-sensitivity and NNS terminator readthrough. The two residues are adjacent in the RNAP II structure (Figure 1). However, the Rpb11-E108G substitution confers a strong flocculation phenotype not seen with Rpb3-K9E, probably due to increased expression of the *FLO1* gene (Chen et al. 2017). Also, the charge reversal substitution Rpb11-E108K is recessive lethal (Steinmetz et al. 2006a), unlike Rpb3-K9E. It is not surprising that substitutions in this region would be pleiotropic, since it is also an interaction site for the Mediator (Soutourina et al. 2011, Schilbach et al. 2023). Further studies will be required to sort out the complex effects of these RNAP II substitutions on gene expression.

## MATERIALS AND METHODS

### Oligonucleotides

See Supplementary Table S4.

### Plasmid construction

pRS314-RPB3 was made by amplifying the *RPB3* gene from yeast strain 46alpha using PCR primers RPB3-UP and RPB3-DOWN, digesting the amplicon at endogenous SacI and PstI sites, and ligating into SacI/PstI digested pRS314. The resulting insert contains 556 base pairs upstream and 425 base pairs downstream of the *RPB3* protein-coding region. pRS306-RPB3, pRS316-RPB3 and pRS317-RPB3 were made by digesting pRS314-RPB3 with SacI and EcoRI and ligating the insert into the SacI/EcoRI-digested plasmid. pRS306-RPB3-TAA and pRS317-RPB3-TAA were made by mutating the *RPB3* stop codon from TAG to TAA by inverse PCR to allow use of the TAG codon for unnatural amino acid incorporation. pRS313-HRP1(RI) was made from DNA amplified from yeast strain 46alpha as described in Goguen and Brow, 2023. pRS315-HRP1, pRS306-HRP1, pRS316-HRP1-L, and pRS315-GFP-HRP1 were made by subcloning the insert from pRS313-HRP1(RI) or pRS313-GFP-HRP1 into the plasmids using BamHI and XhoI sites.

Mutations were made in plasmid-borne *HRP1*, *RPB3*, and *RPB11* by QuikChange or inverse PCR and confirmed by Sanger sequencing. Mutated alleles were either subcloned into an unmutated plasmid backbone or subjected to whole-plasmid sequencing to confirm the absence of unintended mutations. Additional mutations in *HRP1* were acquired in the targeted selection. Double mutations (except for E5K-F207L) were separated by cleavage and ligation using either BamHI and NdeI sites or NdeI and SalI sites. N523D was cloned into pRS315-GFP-HRP1 using AatII and SalI sites.

To create pRS314-RPB3Δ-HIS3 the entire *RPB3* coding region and 18 base pairs of upstream sequence in pRS314-RPB3 were replaced with a 1768 base pair BamHI fragment containing the *HIS3* gene. pRS316-RPB3-HRP1 was made by digesting pRS317-RPB3 with SacI and XhoI and pRS313-HRP1 with XhoI and BamHI, followed by ligation into SacI and BamHI digested pRS316.

pGAC24-SNR13, pGAC24-SNR47, and pGAC24-SNR82 were described in Steinmetz et al. 2001, Chen et al. 2014, and Goguen and Brow 2023, respectively. pG1(LEU2)-HRP1-ORF was constructed by replacing the ACT1-CUP1 ORF in pGAC24 (Lesser and Guthrie 1993) with *HRP1* ORF using the NEB HiFi Assembly Master Mix according to the manufacturer’s protocols. pG1(LEU2)-HRP1-ORF-1-28, pG1(LEU2)-HRP1-ORF-1-40, and pG1(LEU2)-HRP1-ORF-1-129 were cloned by inverse PCR of pG1(LEU2)-HRP1-ORF with primers flanking the region to be deleted (Supplementary Table S3).

### Strain construction

EJS300 was made by transforming strain 46alpha (Lesser & Guthrie, 1993) first with pRS316-RPB3 and selecting for uracil prototrophy, and then performing one-step gene disruption with the linear SacI/EcoRI fragment of pRS314-RPB3Δ-HIS3 and selecting for histidine prototrophy. Integration of *HIS3* at the *RPB3* locus was confirmed by PCR and dependence on a plasmid-borne *RPB3* allele for viability. LWY001 and LYY021 (MW413) were made by transforming EJS300 with pRS314-RPB3 or pRS314-rpb3-K9E, respectively, propagating Trp^+^ cells in CSM -trp liquid medium overnight, and plating cells to medium containing 0.75 mg/mL 5-fluoroorotic acid (5-FOA) to select for loss of pRS316-RPB3. MW445 was made by performing pop-in pop-out protocol by first transforming HindIII linearized pRS306-Hrp1-R317G into MW413, propagating Ura^+^ cells in yeast extract/peptone/dextrose (YEPD) media for 2 passages and then plating cells to medium containing 0.75 mg/mL 5-fluoroorotic acid (5-FOA) to select for pop-outs. The resulting MW445 strain was then verified by genomic DNA extraction, PCR amplification of the HRP1 locus, and Sanger sequencing of the PCR amplicon.

MW402 was made by transforming EJS300 with pRS317-RPB3, propagating Lys^+^ cells in CSM -lys liquid medium overnight, and plating cells to medium containing 0.75 mg/mL 5-fluoroorotic acid (5-FOA) to select for loss of pRS316-RPB3. MW500 was made by transforming MW402 with pRS316-RPB3-HRP1, propagating in CSM -ura liquid medium for 5 passages, and streaking to CSM -lys plate to identify clones that had lost the pRS317-Rpb3 plasmid. One-step gene disruption was then performed on Lys^−^ clones by transformation with *hrp1Δ::KanMX4* amplified from ECG004 (Goguen and Brow 2023) using primers HRP1-Genomic-Upstream and HRP1-Genomic-Downstream and plating to YEPD media containing 0.5 mg/mL G418 disulfate (Alfa Aesar) to select for KanMX4 integration. The disruption was verified by PCR with primers specific for the genomic locus (HRP1-Genomic-Upstream and HRP1-Genomic-Downstream) and Sanger sequencing of the amplicon.

MW403 was made by transforming EJS300 with pRS317-rpb3-K9E, propagating Lys^+^ cells in CSM -lys liquid medium overnight, and plating cells to medium containing 0.75 mg/mL 5-fluoroorotic acid (5-FOA) to select for loss of pRS316-RPB3. MW001 was made by performing modified pop-in pop-out protocol as described in Widlund and Davis, 2004 by first transforming XbaI linearized pRS306-Rpb3-K9E-TAA into MW403, propagating Ura^+^ cells in CSM -ura liquid medium for 6 passages and streaked on CSM -lys plate to confirm the loss of pRS317-rpb3-K9E plasmid. The transformants were then propagated in YEPD media for 2 passages and plated onto medium containing 0.75 mg/mL 5-fluoroorotic acid (5-FOA) to select for pop-outs. The resulting MW001 strain was then verified by genomic DNA extraction, PCR amplification of the RPB3 locus, and Sanger sequencing of the PCR amplicon.

MW507 was made by transforming MW500 with pRS314-rpb3-K9E and pRS315-HRP1, propagating Trp^+^ Leu^+^ cells in CSM -trp-leu liquid medium overnight, and plating cells to medium containing 0.75 mg/mL 5-fluoroorotic acid (5-FOA) to select for loss of pRS316-RPB3-HRP1. MW108 was made by performing modified pop-in pop-out protocol as described in Widlund and Davis, 2004 by first transforming XbaI linearized pRS306-Rpb3-K9E-TAA vector into MW507, propagating Ura^+^ cells in CSM -ura liquid medium for 6 passages and streaked on CSM -trp plate to confirm the loss of pRS314-rpb3-K9E plasmid. The transformants were then propagated in YEPD media for 2 passages and plated onto medium containing 0.75 mg/mL 5-fluoroorotic acid (5-FOA) to select for pop-outs. MW102 was made by transforming pRS316-Hrp1-L into MW108, propagating Ura^+^ cells in CSM -ura liquid medium for 6 passages and streaked on CSM -leu plate to confirm the loss of pRS315-Hrp1. The resulting MW102 was verified by genomic DNA extraction, PCR amplification of the *RPB3* locus, and Sanger sequencing of the PCR amplicon. CSM -leu (1005-010), -trp (1007-010), -his (1006-010), -ura (1004-010), and - leu-trp (1012-010) supplements are from Sunrise Scientific and CSM -lys (Y1896-20G) supplement is from Sigma.

### Serial dilution growth assays

Serial dilutions were conducted in 1:8 dilution series with two biological replicates. Readthrough serial dilutions were stamped onto media lacking leucine and containing various concentrations of CuSO_4_ by a 48-pin inoculation manifold and incubated at 30 °C for 3 or 4 days. Temperature sensitivity serial dilutions were stamped onto rich media with and without supplement adenine (40 mg/L) and incubated at 16 °C (9 days), 23 °C (3 days), 30°C, and 37 °C (2 days). No significant difference was seen with adenine supplementation and only the YEPD results are shown.

### Selection for spontaneous suppressors of the cold-sensitivity of *rpb3-K9E*

Two colonies of LWY021 (labeled A and B) were grown overnight in rich medium (YEPD) at 30 °C and streaked to YEPD plates at 30 °C. Ten colonies from each plate were used to inoculate individual 5 mL YEPD cultures. After two days of growth at 30 °C, 1 mL of each saturated culture was pelleted, 0.8 mL of supernatant was discarded, and the remainder was spread onto a YEPD plate, and the plates incubated at 16 °C. Eight colonies were picked from six plates after 58 days and an additional 12 from 11 plates after 78 days of growth (Table 1).

### Targeted and whole-genome DNA sequencing

Genomic DNA was extracted from 8 mL of saturated YEPD culture using the Thermo Scientific yeast DNA extraction kit (cat. # 78879) with the addition of a 10 min, 37 °C incubation of the isopropanol pellet with 2 mg RNase A in 50 uL water, followed by ethanol precipitation in 1.7 M sodium acetate. The *RPB3* locus from each of 20 putative *rpb3-K9E*-suppressor strains was PCR amplified with oligos RPB3-halfup and pRS MCS downstream or RPB3-halfdown and pRS MCS upstream, and the samples were submitted to Genewiz/Azenta for Sanger sequencing with the primers RPB3-halfup and RPB3-halfdown. All 20 strains had only the *rpb3-K9E* substitution (c.25A>G) and thus are candidates for having obtained extragenic suppressor mutations. The *RPB11* ORF of all 20 strains was also PCR/Sanger sequenced with oligos RPB11-F and RPB11-R and found to have no mutations. For AmpliSeq targeting of transcription and chromatin related loci, genomic DNA from candidates and parental strains were processed with the AmpliSeq for Illumina Custom DNA Panel kit deploying a panel of 1917 primer pairs targeting 310 genes similar to a previously deployed approach (Brow 2019). The panel of 310 genes was designed to encompass the open reading frames of transcription machinery and transcription-associated genes (Sikorski et al. 2011) by 1917 overlapping amplicons ranging from 107-331bps. AmpliSeq libraries were generated and sequenced by 2x 250bp cycle paired-end Illumina MiSeq sequencing at the Texas A&M University Institute for Genome Sciences and Society core.

For whole genome sequencing, 0.5 ug of genomic DNA from each yeast strain, prepared as described above, was sheared to a target length of 300 bp using a Covaris S220 with the microTUBE-130 AFA. Illumina sequencing libraries were prepared using the NEBNext Ultra II DNA Library Prep Kit (catalog # E7645S) and purified using Beckman Coulter AMPure XP beads (catalog # A63881). Libraries were sequenced on an Illumina NextSeq 1000 with a 100-cycle P2 flow cell (2×50bp, Illumina Cat. # 20046811). The total reads for each sample ranged from 6.6 to 16.9 million and the fraction of aligned reads ranged from 93.0 % to 98.8 %, with final coverage of 27- to 69-fold.

### Informatics analysis of sequencing results

For mutation identification by AmpliSeq, sequencing reads were first stripped of adaptor sequences and amplicon amplifying oligos. We used cutadapt version 2.10 (Martin 2011) to strip reads of 3’ adaptor sequence followed by trimming 3’ non-internal oligos in the adaptor-trimmed reads. All reads were then trimmed at the 5’ of 6 bases representative of amplicon 5’ oligo trimming. Adaptor and terminal oligo-free reads were then processed by MutantHuntWGS v1.1 (Ellison et al. 2020), a variant calling package for *Saccharomyces cerevisiae* with analysis pipeline customizable through user-defined flag inputs for preloaded packages with defined versions. Using MutantHuntWGS v1.1 paired-end fastq reads (MutanthuntWGS v1.1 flag “-r paired”) were aligned to *Saccharomyces cerevisiae* S288C reference genome R64-2-1 by bowtie2 version 2.2.9 (Langmead and Salzberg 2012). Resulting bam files were sorted and indexed by SAMtools version 1.3.1 (Li et al. 2009), followed by genotype likelihood calculations. Variants were called by BCFtools version 1.3.1 (Danecek et al. 2021) using a haploid ploidy file (MutanthuntWGS v1.1 flag “-p ploidy_n1.txt”), followed by differential variant calling of mutant vcf files relative to parental vcf files by VCFtools version 0.1.14 (Danecek et al. 2011) and variants filtered using a variant score cutoff of 30 (MutanthuntWGS v1.1 flag “-s 30”). Mutants were then annotated using SNPeff version 4.3p (Cingolani et al. 2012) and SIFT4G (Vaser et al. 2016), the latter using the EF4.74 *Saccharomyces cerevisiae* library for assessing amino acid change impact.

For whole-genome sequencing, variants were identified using *breseq* v0.36.1 with default parameters (Deatherage and Barrick, 2014) by comparing to S288C reference genome R64-3-1_202110421. Because 46alpha is not closely related to S288C, there were 4779 variants common to all 20 strains and an additional 182 present in 19 or 18 strains. Likely causal variants were identified for 18 strains (Table 1) by manually inspecting the 172 variants present in 1-5 strains.

### Targeted selection for suppressors of *rpb3-K9E* in *HRP1*

The *HRP1* ORF was randomized using Taq polymerase as described in Goguen and Brow (2023). *HRP1* ORF Gapped pRS315-HRP1 backbone was amplified using primers HRP1-Gapped-F and HRP1-Gapped-R followed by DpnI digest and cleaned up using the GeneJET Gel Extraction Kit (Thermo Scientific, Cat. # K0691). The randomized *HRP1* ORF and gapped pRS315-HRP1 backbone were cotransformed into MW102 and plated onto a synthetic dropout media lacking leucine. Transformation plates were replica plated onto synthetic complete media containing 0.75 mg/mL 5-fluoroorotic acid (5-FOA) to select for loss of the pRS316-HRP1 shuffle plasmid. The 5-FOA plates were replica plated onto YEPD medium and incubated at 16°C for 5 or 6 days. Plasmid were rescued from cold-resistant colonies using the GeneJET Plasmid Miniprep Kit (Thermo Scientific, Cat. # K0502) following cell lysis by vortexing with glass beads (425–600 μm, Sigma Aldrich, Cat. # G8771-100G). The rescued pRS315-HRP1 plasmids were transformed into DH5α, harvested using the GeneJET Plasmid Miniprep Kit, and whole plasmid sequenced to identify all mutations.

### Western blots

Western blots were done as described in Goguen and Brow 2023 with the following changes. Yeast cells were harvested at OD_600_ of 0.75-1.25. Protein Gels (Bio-Rad, Cat. # 4561086) were run at 150 V for 56 min and electroblotted onto Immobilon-P PVDF membrane (Millipore, Cat. # IPVH00010) at 95 V, 4 °C, for 60 min. After blocking, blots were incubated with GFP polyclonal antibody (1:3,000, Invitrogen, Cat. # A11122) at 4 °C overnight, and then with HRP-conjugated goat anti rabbit IgG (1:5,000, Pierce Scientific, Cat. # 31460) at 4 °C for 120 min. Bands were quantified using ICY.

### Fluorescence Imaging

Cells were grown overnight in CSM -leu to OD_600_ of 0.9-1.1 and directly dropped onto microscope slides. Fluorescence images were taken using AxioPlan 2 Microscope (Zeiss) and processed using ICY.

## ACKNOWLEDGEMENTS

We thank Eric Steinmetz for creating the EJS300 yeast strain, Lynn Weaver for creating yeast strains LWY001 and LWY021 and initial analysis of *RPB3* alleles, Jacob Eckmann, Kevin Myers, and Patricia Kiley for whole-genome sequencing and variant analysis, and Michael Henry for Hrp1 anti-serum. This work was supported by NIH grants R01 GM082956 and R35 GM118075 to DAB and by R01 GM120450 and R35 GM144116 to CDK.

---

**Table S1.** Yeast strains used in this study.

| **Strain** | **Genotype** | **Source** |
| --- | --- | --- |
| 46alpha | *MATalpha cup1Δ::ura3 ura3-52 his3-Δ200 trp1-Δ63 lys2-801 ade2-1 leu2-Δ1* | Lesser and Guthrie, 1993 |
| EJS300 | *MATalpha rpb3Δ::HIS3 cup1Δ::ura3 ura3-52 his3-Δ200 trp1-Δ63 lys2-801 ade2-1 leu2-Δ1 [pRS316-RPB3]* | This study |
| LWY001 | *EJS300 [pRS314-RPB3]* | This study |
| LWY021 | *EJS300 [pRS314-rpb3-K9E]* | This study |
| MW001 | *MATalpha rpb3-K9E-TAA cup1Δ::ura3 ura3-52 his3-Δ200 trp1-Δ63 lys2-801 ade2-1 leu2-Δ1* | This study |
| MW102 | *MATalpha rpb3-K9E-TAA hrp1Δ::KanMX4 cup1Δ::ura3 ura3-52 his3-Δ200 trp1-Δ63 lys2-801 ade2-1 leu2-Δ1* | This study |
| MW500 | *MATalpha rpb3Δ::HIS3 hrp1Δ::KanMX4 cup1Δ::ura3 ura3-52 his3-Δ200 trp1-Δ63 lys2-801 ade2-1 leu2-Δ1 [pRS316-RPB3-HRP1]* | This study |
| XCY361 | *MATα his3-Δ1 leu2-Δ0 lys2-Δ0 ura3-Δ0 NHP6A-TagRFP-T::nat* | Chen et al., 2014 |
| MW410 | *EJS300 [pRS314-Rpb3]* | This study |
| MW411 | *EJS300 [pRS314-Rpb3-K9A]* | This study |
| MW412 | *EJS300 [pRS314-Rpb3-K9D]* | This study |
| MW413 | *EJS300 [pRS314-Rpb3-K9E]* | This study |
| MW414 | *EJS300 [pRS314-Rpb3-K9M]* | This study |
| MW415 | *EJS300 [pRS314-Rpb3-K9R]* | This study |
| MW416 | *EJS300 [pRS314-Rpb3-K9Q]* | This study |
| MW445 | *EJS300 [pRS314-Rpb3-K9E] hrp1-R317G* | This study |
| MW503 | *MW500 [pRS314-Rpb3; pRS315-Hrp1]* | This study |
| MW504 | *MW500 [pRS314-Rpb3; pRS315-Hrp1-R317G]* | This study |
| MW505 | *MW500 [pRS314-Rpb3; pRS315-Hrp1-R317Q]* | This study |
| MW506 | *MW500 [pRS314-Rpb3; pRS315-Hrp1-R317E]* | This study |
| MW507 | *MW500 [pRS314-Rpb3-K9E; pRS315-Hrp1]* | This study |
| MW508 | *MW500 [pRS314-Rpb3-K9E; pRS315-Hrp1-R317G]* | This study |
| MW509 | *MW500 [pRS314-Rpb3-K9E; pRS315-Hrp1-R317Q]* | This study |
| MW510 | *MW500 [pRS314-Rpb3-K9E; pRS315-Hrp1-R317E]* | This study |
| MW511 | *MW500 [pRS314-Rpb3; pRS315-GFP-Hrp1]* | This study |
| MW513 | *MW500 [pRS314-Rpb3-K9E; pRS315-GFP-Hrp1]* | This study |
| MW515 | *MW500 [pRS314-Rpb3-K9E; pRS315-GFP-Q29X]* | This study |
| MW517 | *MW500 [pRS314-Rpb3-K9E; pRS315-GFP-Q41X]* | This study |
| MW519 | *MW500 [pRS314-Rpb3-K9E; pRS315-GFP-Q130X]* | This study |
| MW521 | *MW500 [pRS314-Rpb3-K9E; pRS315-GFP-N523D]* | This study |
| MW523 | *MW500 [pRS314-Hrp1-Rpb3-K9E; pG1(LEU2)-HRP1-1-28]* | This study |
| MW525 | *MW500 [pRS314-Hrp1-Rpb3-K9E; pG1(LEU2)-HRP1-1-40]* | This study |
| MW527 | *MW500 [pRS314-Rpb3-K9E; pRS315-Hrp1-E5K-F207L]* | This study |
| MW529 | *MW500 [pRS314-Rpb3-K9E; pRS315-Hrp1-F207L]* | This study |
| MW531 | *MW500 [pRS314-Rpb3-K9E; pRS315-Hrp1-Q29X]* | This study |
| MW533 | *MW500 [pRS314-Rpb3-K9E; pRS315-Hrp1-E258G]* | This study |
| MW535 | *MW500 [pRS314-Rpb3-K9E; pRS315-Hrp1-K309T-N457S]* | This study |
| MW537 | *MW500 [pRS314-Rpb3-K9E; pRS315-Hrp1-Q41X]* | This study |
| MW539 | *MW500 [pRS314-Rpb3-K9E; pRS315-Hrp1-Q130X]* | This study |
| MW541 | *MW500 [pRS314-Rpb3-K9E; pRS315-Hrp1-V289A-G412G]* | This study |
| MW543 | *MW500 [pRS314-Rpb3-K9E; pRS315-Hrp1-V185A]* | This study |
| MW545 | *MW500 [pRS314-Rpb3-K9E; pRS315-Hrp1-K379E]* | This study |
| MW547 | *MW500 [pRS314-Rpb3-K9E; pRS315-Hrp1-Q385H]* | This study |
| MW549 | *MW500 [pRS314-Rpb3-K9E; pRS315-Hrp1-T219A-N523D]* | This study |
| MW551 | *MW500 [pRS314-Rpb3-K9E; pRS315-Hrp1-M373K]* | This study |
| MW559 | *MW500 [pRS314-Rpb3-K9E; pRS315-Hrp1-T219A]* | This study |
| MW561 | *MW500 [pRS314-Rpb3-K9E; pRS315-Hrp1-N523D]* | This study |
| MW563 | *MW500 [pRS314-Rpb3-K9E; pRS315-Hrp1-K309T]* | This study |
| MW565 | *MW500 [pRS314-Rpb3-K9E; pRS315-Hrp1-N457S]* | This study |
| MW567 | *MW500 [pRS314-Rpb3-K9E; pRS315-Hrp1-E5K]* | This study |
| MW569 | *MW500 [pRS314-Rpb3; pRS315-Hrp1-E5K]* | This study |
| MW571 | *MW500 [pRS314-Rpb3-K9E; pRS315-Hrp1-Y383H]* | This study |
| MW573 | *MW500 [pRS314-Rpb3-K9E; pRS315-Hrp1-M373K-K379E]* | This study |

**Table S3.** Mutations identified in HRP1 targeted *rpb3-K9E*-suppressor strains.

| Clone | Days at 16° | Strain # | Color | Identified mutation(s) | Identified DNA substitution(s) |
| --- | --- | --- | --- | --- | --- |
| 200B-1 | 5 | MW911 | red | T20T | A60C |
| 200B-2 | 5 | MW912 | white | WT | WT |
| 200B-3 | 5 | MW913 | red |  |  |
| 200B-4 | 5 | MW914 | red | WT | WT |
| 400-1 | 5 | MW921 | red | K309T; N457S | A626C; A1370G |
| 400-2 | 5 | MW922 | white | WT | WT |
| 400-3 | 5 | MW923 | red | Q385H | A1155T |
| 400-4 | 5 | MW924 | red | E258G | A773G |
| 200A-1 | 6 | MW901 | red | V185A | T554C |
| 200A-2 | 6 | MW902 | red | WT | WT |
| 200A-3 | 6 | MW903 | red | WT | WT |
| 200A-4 | 6 | MW904 | red |  |  |
| 200A-5 | 6 | MW905 | red | T219A; N523D | A655G; A1567G |
| 200A-6 | 6 | MW906 | red | Q41X; S60S | C121T; T180C |
| 200A-7 | 6 | MW907 | red | M373K | T1118A |
| 200A-8 | 6 | MW908 | red | V289A; G412G | T866C; G1236A |
| 200B-5 | 6 | MW915 | red | WT | WT |
| 200B-6 | 6 | MW916 | red | WT | WT |
| 200B-7 | 6 | MW917 | red | WT | WT |
| 200B-8 | 6 | MW918 | red | R255R | A765G |
| 200B-9 | 6 | MW919 | red | Q29X | C85T |
| 200B-10 | 6 | MW920 | red | WT | WT |
| 400-5 | 6 | MW925 | red | WT | WT |
| 400-6 | 6 | MW926 | red | F207L | T619C |
| 400-7 | 6 | MW927 | red | Q41X | C121T |
| 400-8 | 6 | MW928 | red | Q130X | C388T |
| 400-9 | 6 | MW929 | red | E5K; F207L | G13A; T619C |
| 400-10 | 6 | MW930 | red | K379E | A1134G |

**Table S4.**
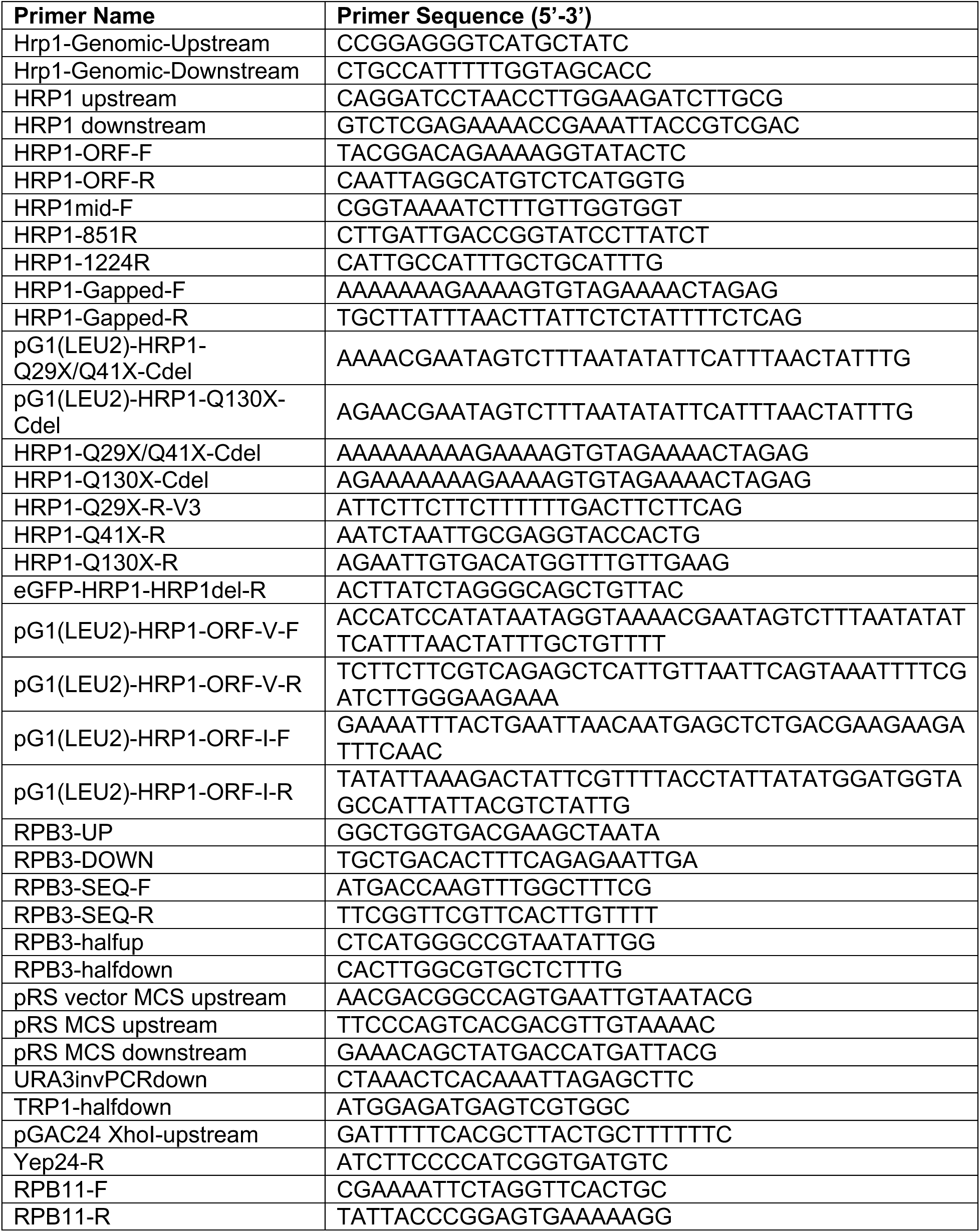
Oligonucleotides used in this study.

**Figure S1.**
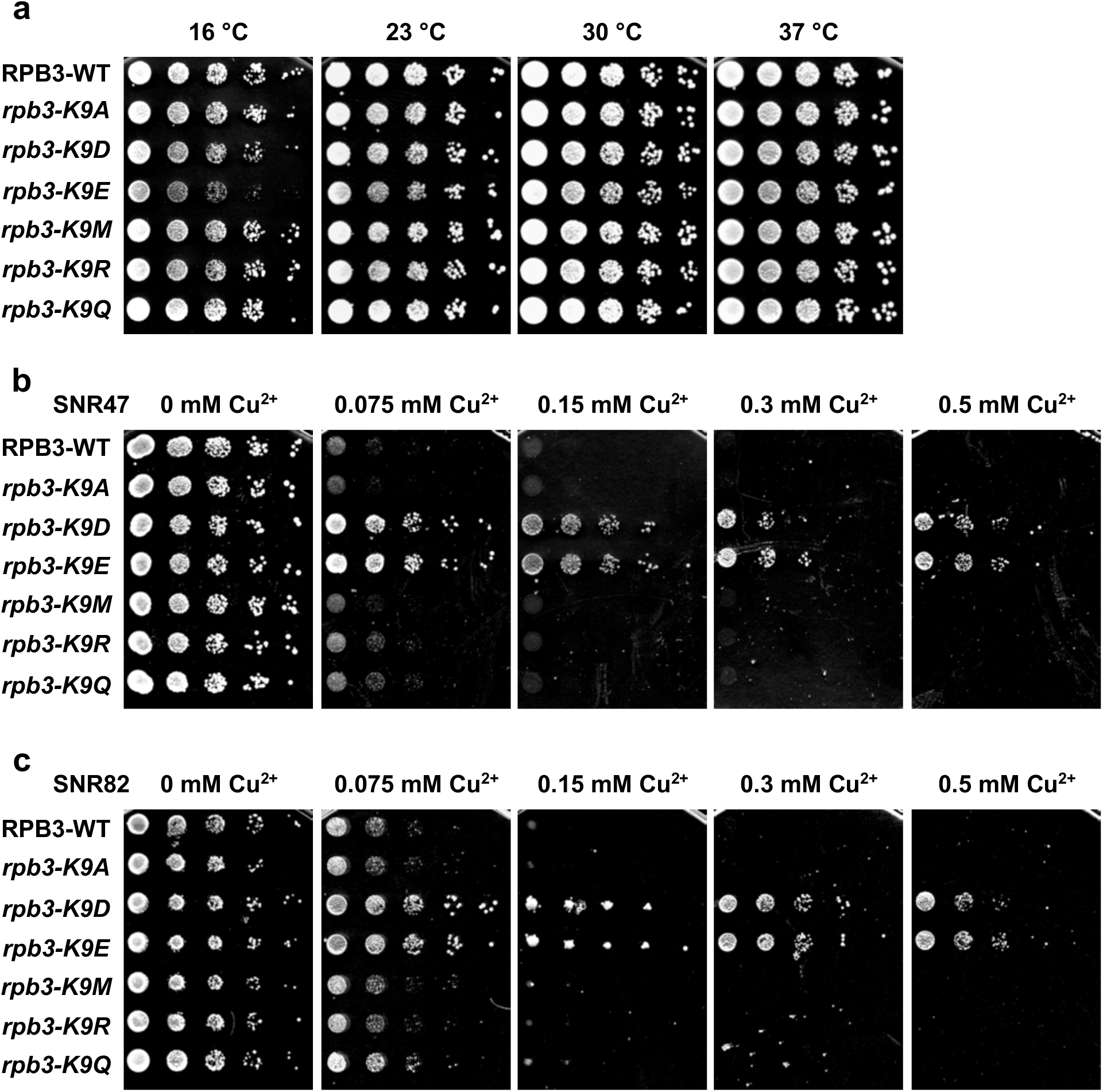
Biological replicates of data shown in Figure 2. **a)** Serial 8-fold dilutions of a haploid *Δrpb3* strain containing the indicated alleles of *RPB3* on a centromere plasmid were spotted on YEPD plates and incubated at the indicated temperatures for two (30, 37 °C), three (23 °C), or nine (16 °C) days. Serial 8-fold dilutions of haploid strains containing the indicated alleles of *RPB3* and the (**b**) *SNR47* or (**c**) *SNR82* ACT1-CUP1 reporter on separate centromere plasmids were spotted on synthetic complete medium containing the indicated concentration of copper sulfate and incubated at 30 °C for 4 days.

**Figure S2.**
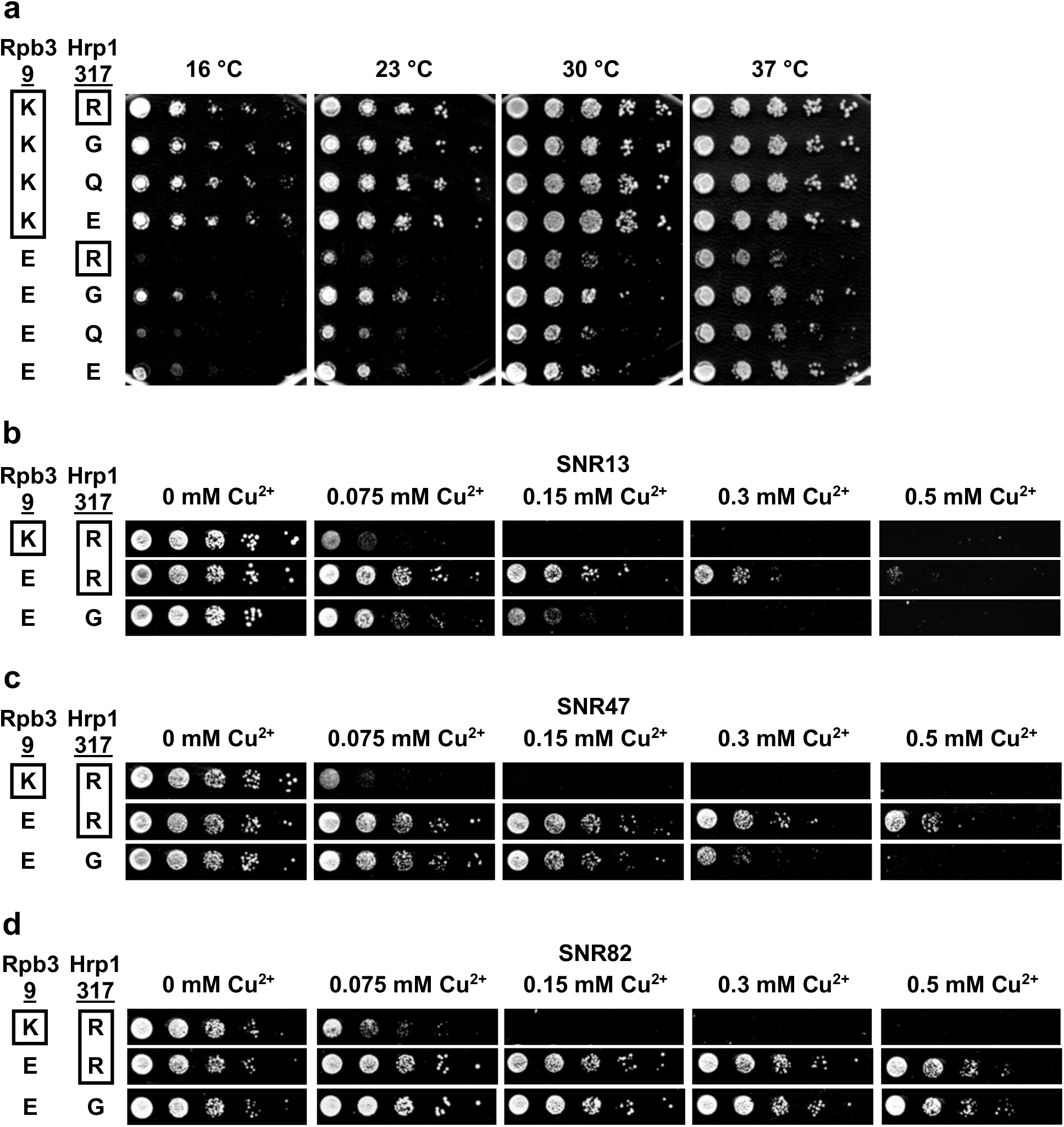
Biological replicates of data shown in Figure 4. **a)** Serial dilutions of haploid strains containing the indicated alleles of *RPB3* and *HRP1* on separate centromere plasmids were spotted onto YEPD plates and incubated at the indicated temperatures for 2 (30, 37 °C), 3 (23 °C), and 7 (16 °C) days. Wildtype residues are boxed. **b-d**) Serial dilutions of haploid strains containing the indicated alleles of *RPB3* on a centromere plasmid with indicated alleles of *HRP1* in the genome on CSM -Leu medium containing the indicated concentrations of copper sulfate were incubated at 30 °C for 4 days. All rows for each panel came from the same plate.

**Figure S3.**
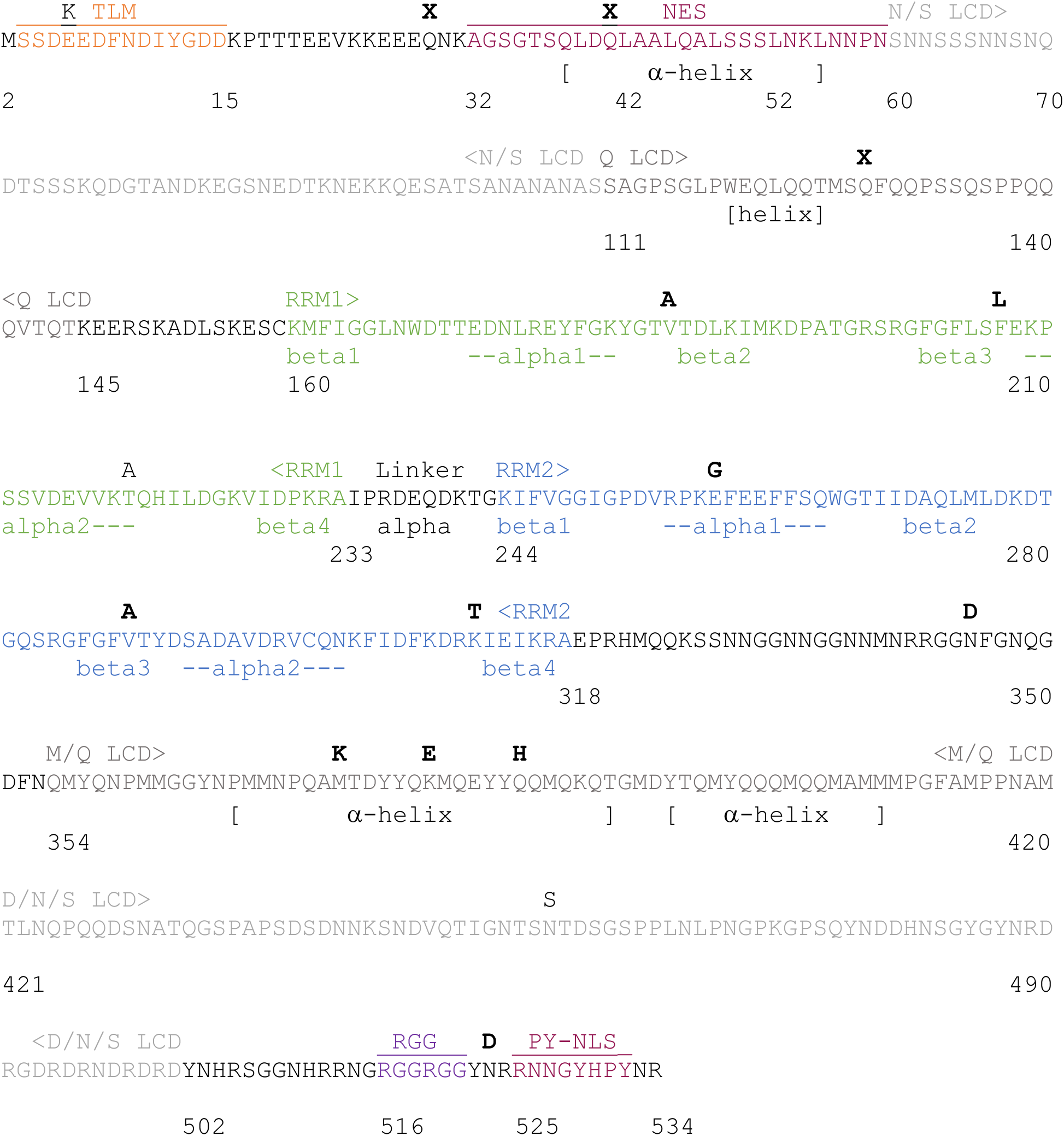
Annotated Hrp1 sequence with domains highlighted. Alpha-fold predicted secondary structure elements in the LCDs are indicated with [ ]. Locations of alpha-helices and beta-strands in the RRMs are based on the NMR structure (Pérez-Cañadillas 2006). *rpb3-*K9E-suppressor substitutions are shown above the sequence in bold. The unbolded substitutions are “passengers” obtained with suppressors. Adapted from Goguen and Brow, 2023.

**Figure S4.**
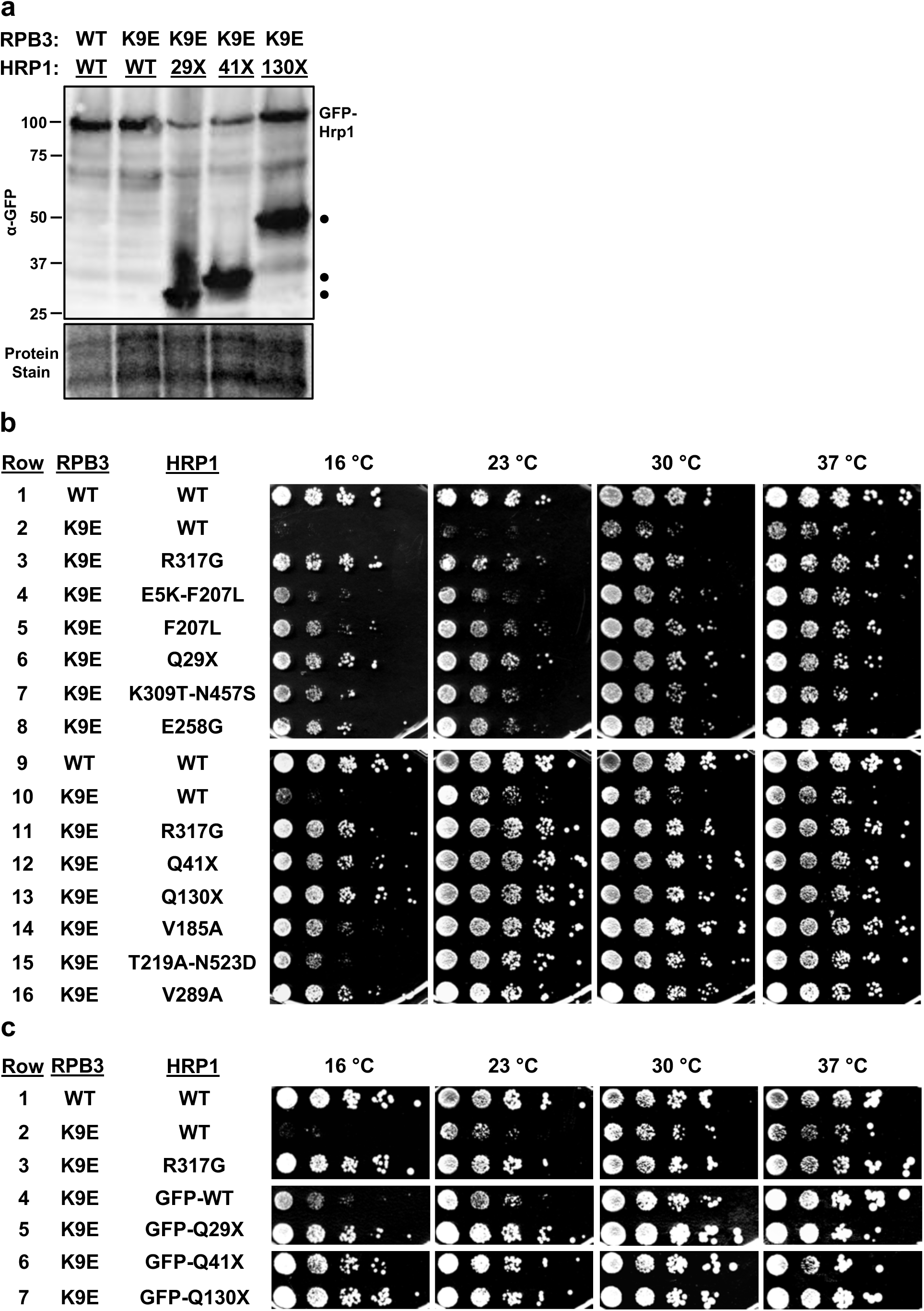
Biological replicates of data shown in Figure 5. **a)** Anti-GFP Western Blot of cell extracts from *RPB3*-*HRP1* double disruption strains with the indicated mutations in *RPB3* and *HRP1* on separate centromere plasmids. A duplicate Coomassie blue stained gel, part of which is shown below, was used to normalize for total protein loaded. **b,c**) Serial dilutions (1:8) of haploid strains containing the indicated alleles of *RPB3* and *HRP1* on separate centromere plasmids were spotted onto YEPD plates and incubated at the indicated temperatures for 2 (30, 37 °C), 3 (23 °C), or 9 (16 °C) days.

**Figure S5.**
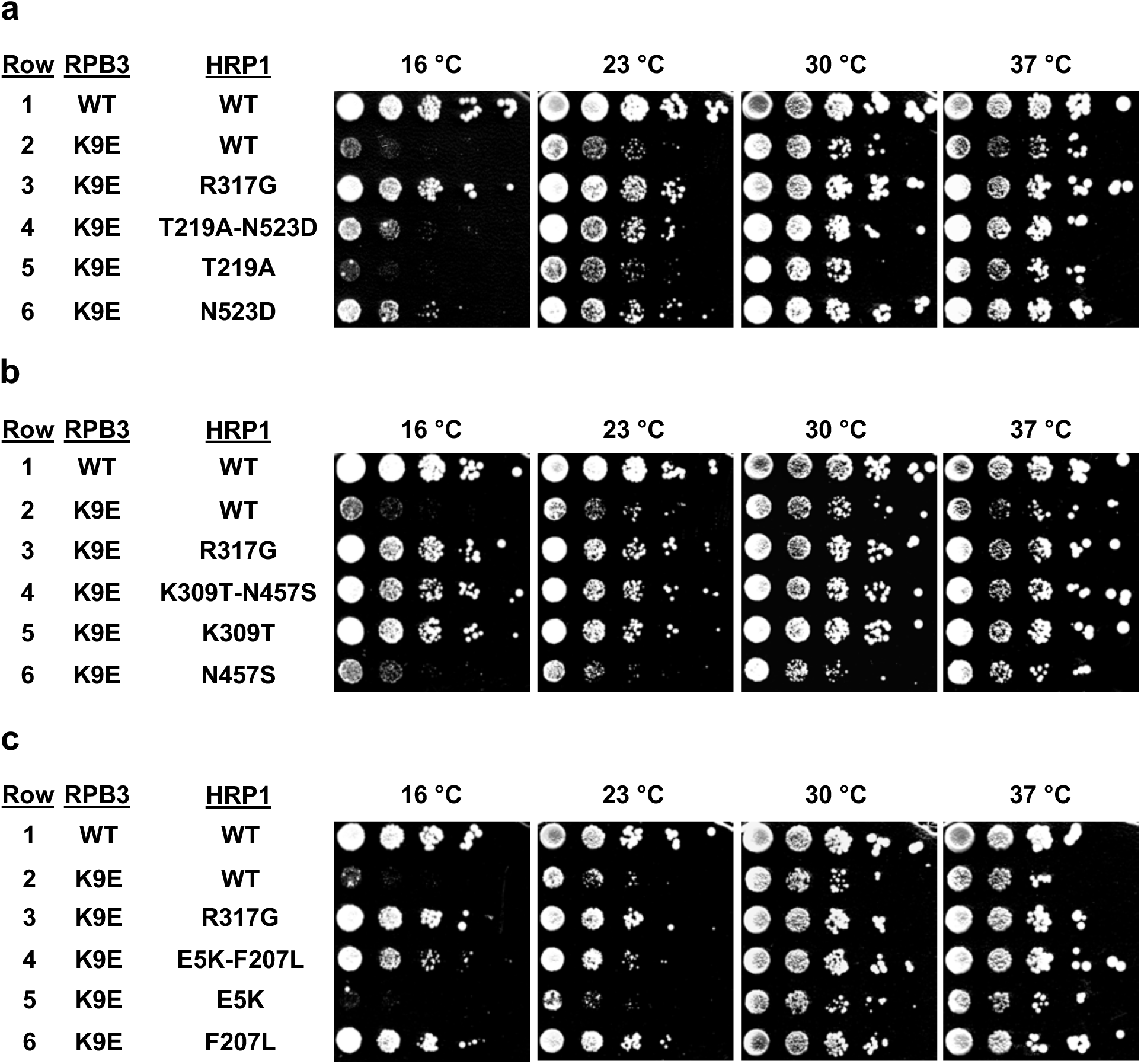
Biological replicates of data in Figure 6. Serial dilutions (1:8) of haploid strains containing the indicated alleles of *RPB3* and *HRP1* on separate centromere plasmids were spotted onto YEPD plates and incubated at the indicated temperatures for 2 (30, 37 °C), 3 (23 °C), and 9 (16 °C) days. The starting double mutant is in row 4 of each panel and each single mutant is in rows 5 and 6.

**Figure S6.**
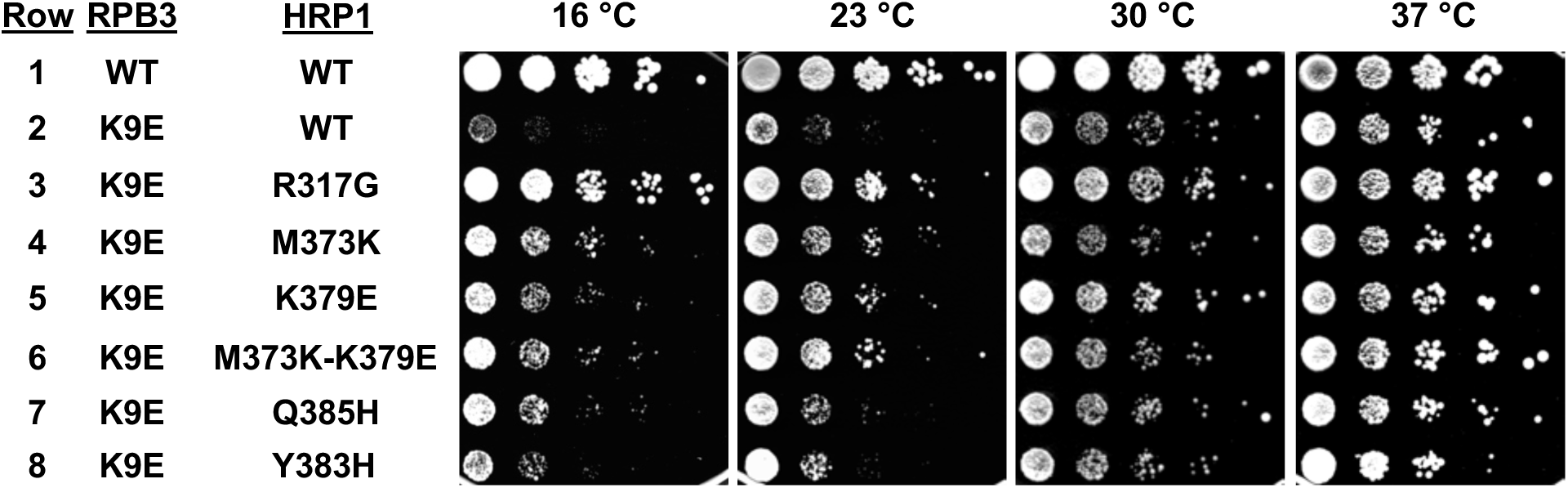
Biological replicates of data in Figure 7. Serial dilutions (1:8) of haploid strains containing the indicated alleles of *RPB3* and *HRP1* on separate centromere plasmids were spotted onto YEPD plates and incubated at the indicated temperatures for 2 (30, 37 °C), 3 (23 °C), and 9 (16 °C) days.

**Figure S7.**
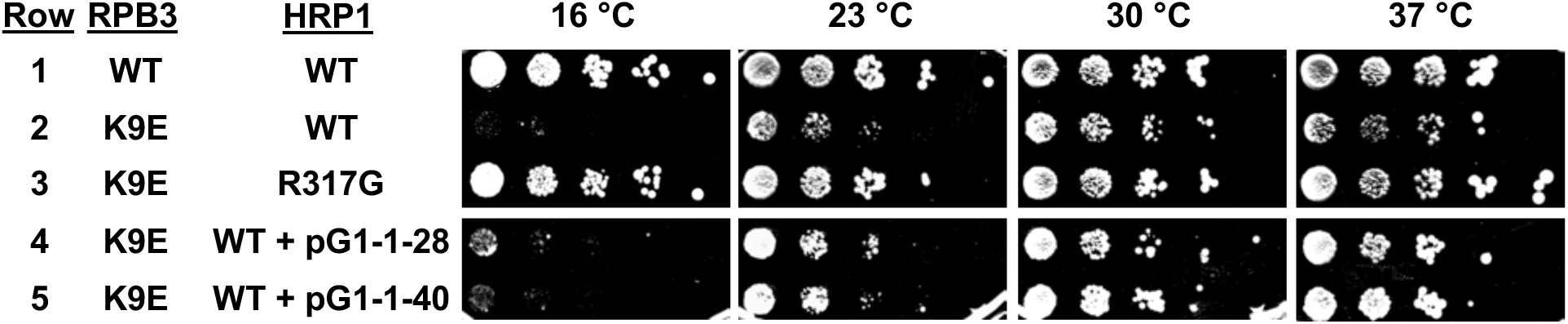
Biological replicates of data in Figure 8. Serial dilutions (1:8) of haploid strains containing the indicated alleles of *RPB3* and *HRP1* on separate centromere plasmids (Row 1-3) or on the same centromere plasmid with Hrp1 overexpression construct (Row 4-5) were spotted onto YEPD plates and incubated at the indicated temperatures for 2 (30, 37 °C), 3 (23 °C), and 9 (16 °C) days.

**Figure S8.**
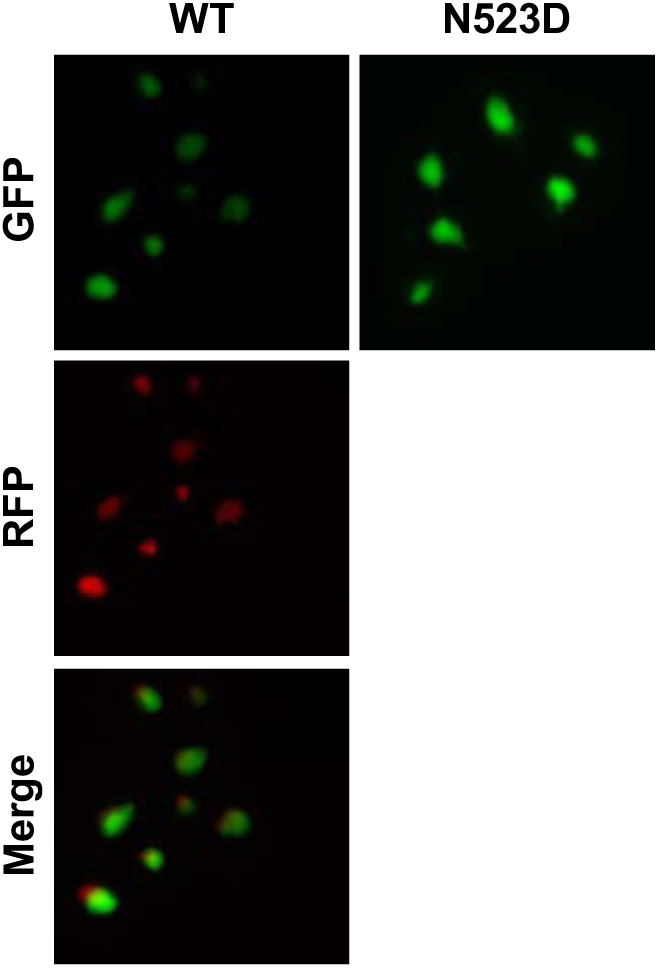
Hrp1-N523D does not impact Hrp1 nuclear localization. GFP tagged Hrp1-WT and Hrp1-N523D on centromere plasmid were transformed into cells with integrated RFP tagged NHP6A nuclear marker. Cells were analyzed by fluorescence imaging after growth to 1.0 OD.

**Figure S9.**
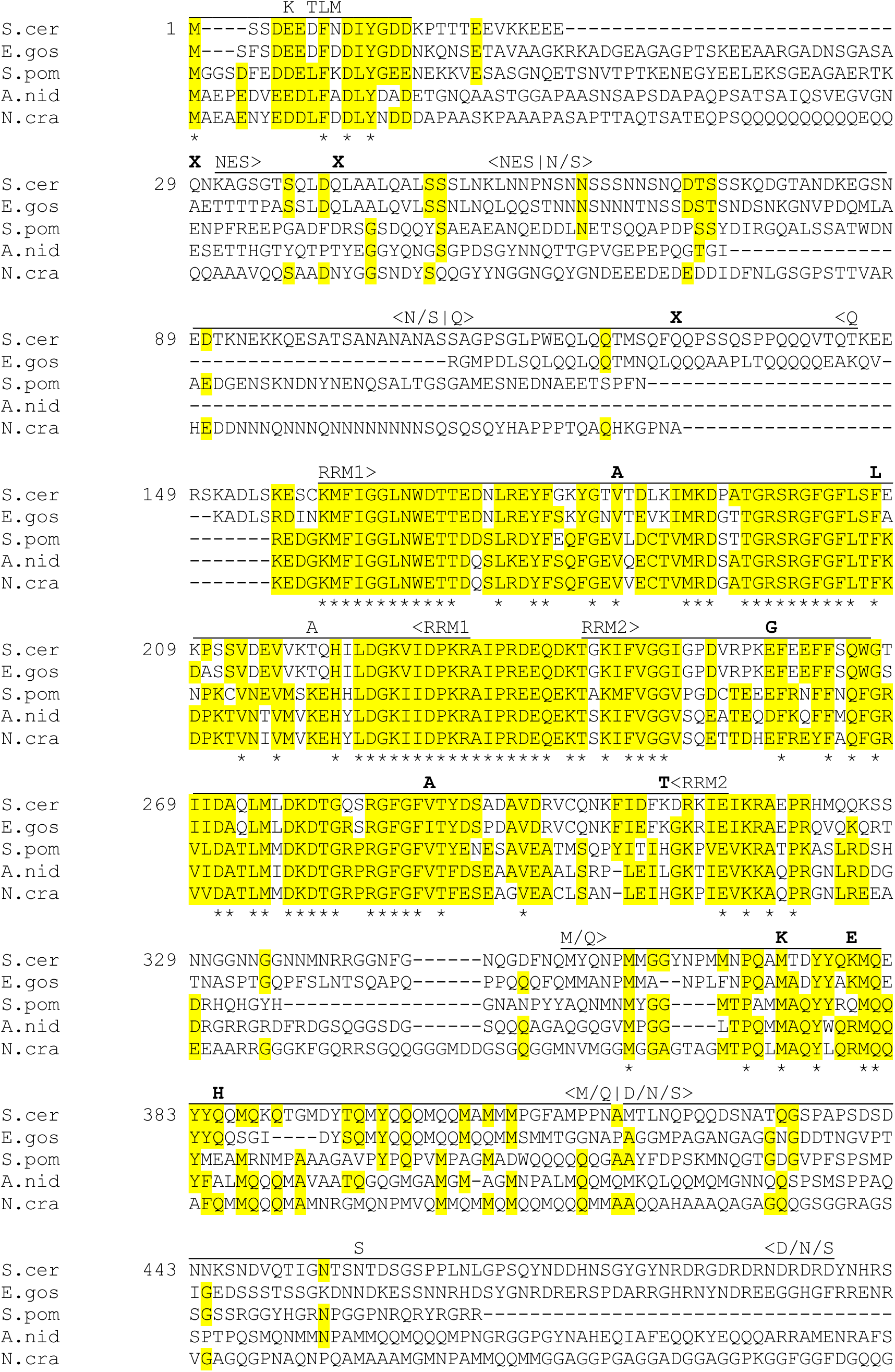
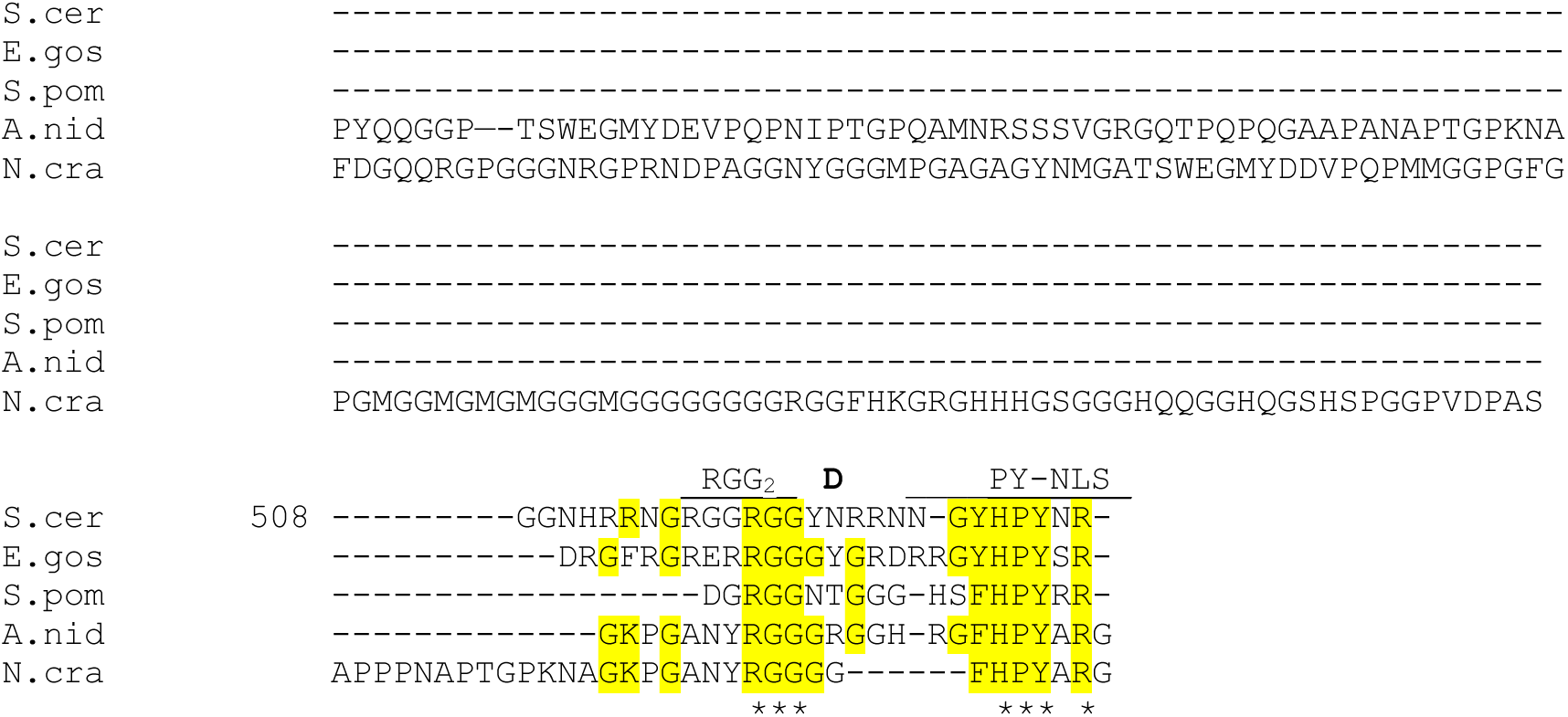
Hrp1 fungal sequence alignment. Highlighted regions are conserved (identical or similar amino acids) in at least three out of five species. Residues that are identical in all five species are indicated with (*). *rpb3-*K9E-suppressor substitutions are shown in bold above the sequences. Three “passenger” substitutions in double mutant alleles are shown in unbolded font. *Saccharomyces cerevisiae* (S.cer)*, Eremothecium gossypii (E.gos), Schizosaccharomyces pombe (S.pom), Aspergillus nidulans (A.nid), and Neurospora crassa (N.cra)*. Adapted from Goguen and Brow, 2023.

**Figure S10.**
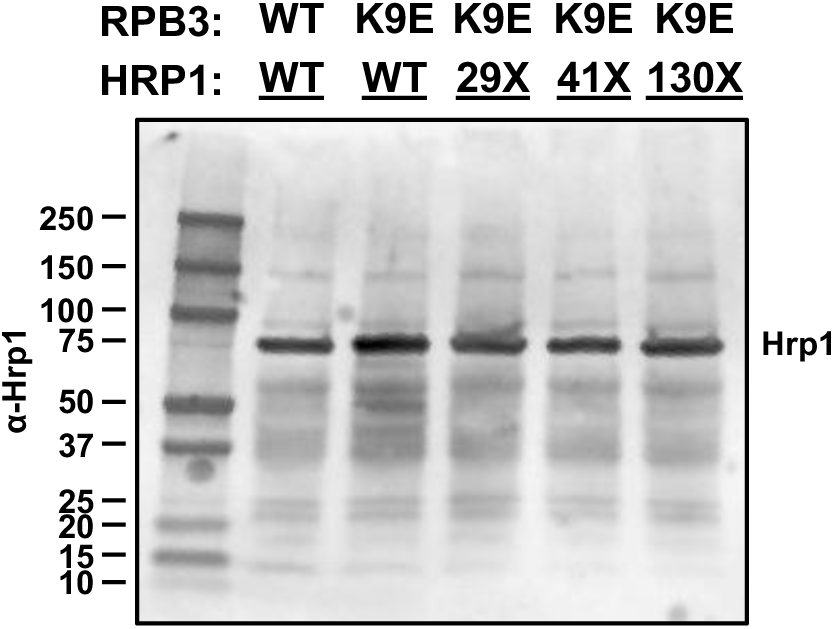
Anti-Hrp1 polyclonal antibodies do not detect truncated Hrp1 proteins. Anti-Hrp1 Western Blot of cell extracts from *RPB3*-*HRP1* double disruption strains with the indicated mutations in *RPB3* and *HRP1* on separate centromere plasmids.

